# The nuclear periphery confers repression on H3K9me2-marked genes and transposons to shape cell fate

**DOI:** 10.1101/2024.07.08.602542

**Authors:** Harold Marin, Eric Simental, Charlie Allen, Eric Martin, Barbara Panning, Bassem Al-Sady, Abigail Buchwalter

## Abstract

Heterochromatic loci marked by histone H3 lysine 9 dimethylation (H3K9me2) are enriched at the nuclear periphery in metazoans, but the effect of spatial position on heterochromatin function has not been defined. Here, we remove three nuclear lamins and lamin B receptor (LBR) in mouse embryonic stem cells (mESCs) and show that heterochromatin detaches from the nuclear periphery. Mutant mESCs sustain naïve pluripotency and maintain H3K9me2 across the genome but cannot repress H3K9me2-marked genes or transposons. Further, mutant cells fail to differentiate into epiblast-like cells (EpiLCs), a transition that requires the expansion of H3K9me2 across the genome. Mutant EpiLCs can silence naïve pluripotency genes and activate epiblast-stage genes. However, H3K9me2 cannot repress markers of alternative fates, including primitive endoderm. We conclude that the nuclear periphery controls the spatial position, dynamic remodeling, and repressive capacity of H3K9me2-marked heterochromatin to shape cell fate decisions.

## Introduction

Eukaryotic genomes are organized into compartments of transcriptionally active euchromatin and silent heterochromatin. Heterochromatin shapes cell fate decisions by restricting gene expression and safeguards genome integrity by inhibiting transposon activity^1^. Across eukaryotes, heterochromatin is abundant at the nuclear periphery while euchromatin resides in the nuclear interior^2–5^. In mammals, the peripheral position of heterochromatin is established in embryonic nuclei shortly after fertilization and before other features of genome folding, such as topologically associating domains, appear^6^. The nuclear periphery appears to have an intrinsic and conserved capacity to repress transcription. In *S. pombe,* heterochromatin recruitment to the nuclear periphery enables the long-term epigenetic memory of cell state by antagonizing nucleosome turnover^7^. In mammals, artificial tethering of loci to the nuclear periphery induces silencing^8–10^. Association to the nuclear periphery and transcriptional repression are strongly correlated, and many lineage-irrelevant genes are recruited to the nuclear periphery during differentiation processes^11,12^. These observations have led to the prevailing model that peripheral heterochromatin positioning promotes the establishment of cell fate by repressing alternative fate genes. However, this model has not yet been tested, and the mechanism by which the nuclear periphery confers repression on associated chromatin remains unknown.

Recently, it has become clear that histone H3 lysine 9 di- and/or tri-methylation (H3K9me2/3) is uniquely enriched on peripheral heterochromatin in many eukaryotes^7,13–16^ and is required for the formation of this compartment^16–18^. H3K9me2 is predominantly deposited by the heterodimeric G9a/GLP enzyme (also known as EHMT2 and EHMT1)^19,20^. H3K9me2 promotes the repression of both genes and transposons^19,21–25^ and is essential for mammalian pre-implantation development^19,20,24^. However, H3K9me2-modified chromatin is permissive to transcription in some contexts^26–28^, which implies that H3K9me2 alone is not sufficient to induce repression.

The receptor proteins that tether H3K9me2/3-marked loci to the nuclear periphery vary across eukaryotes and include components of the inner nuclear membrane (INM) and of the nuclear lamina, a meshwork formed by lamin proteins that underlies the INM. For instance, the INM protein Amo1 tethers H3K9me3 to the nuclear periphery in *S. pombe*^7^. In *C elegans,* the INM protein CEC-4 tethers H3K9me2/3-marked chromatin in embryos but is dispensable for this process in differentiated tissues, perhaps due to compensation by other proteins^13,29^. In mammals, the lamin isoform lamin A/C and the nuclear membrane protein lamin B receptor (LBR) each contribute to the spatial position of heterochromatin^30^, although it is unknown whether they recognize H3K9me2/3. H3K9me2 is highly enriched in lamina-associated domains (LADs) of chromatin in mammals^14,18,31–33^, raising the possibility that the nuclear lamins and/or LBR control the peripheral positioning of heterochromatin bearing this modification.

Here, we disrupt the lamins and LBR in mouse embryonic stem cells (mESCs) to dissect the functions of heterochromatin positioning during early mammalian development. We show that these proteins control the spatial positioning of heterochromatin and discover that H3K9me2 is unable to effectively repress either transposons or lineage-specific genes when displaced from the nuclear periphery, indicating that heterochromatin positioning enhances the repression of H3K9me2-modified chromatin. Finally, we show that displacing heterochromatin from the nuclear periphery impairs the timely restriction of gene expression that shapes cell fate during differentiation. Altogether, this work reveals how the nuclear periphery sculpts the function of heterochromatin.

## Results

### The lamins and LBR recruit heterochromatin to the nuclear periphery in pluripotent cells

A strong correlation between LADs and H3K9me2-marked chromatin has been reported in a range of mammalian cell types^14,33^. However, mESCs lacking all lamin isoforms are viable and pluripotent, retain peripheral heterochromatin positioning, and exhibit modest changes to genome folding and gene expression^34–36^. We noted that both wild type (WT) and lamin triple knockout (TKO) mESCs cultured in naïve conditions (2i + LIF) express high levels of LBR which remains enriched at the nuclear periphery even in the absence of the lamin proteins in TKO mESCs (Fig. S1B), while H3K9me2-marked chromatin also remains enriched at the nuclear periphery in both WT and TKO mESCs (Fig. 1A-B). These observations led us to hypothesize that LBR sustains heterochromatin organization in this context^30^, and to propose that lamin TKO mESCs are a useful sensitized system in which to dissect the functions of heterochromatin positioning in mammals.

**Figure 1.**
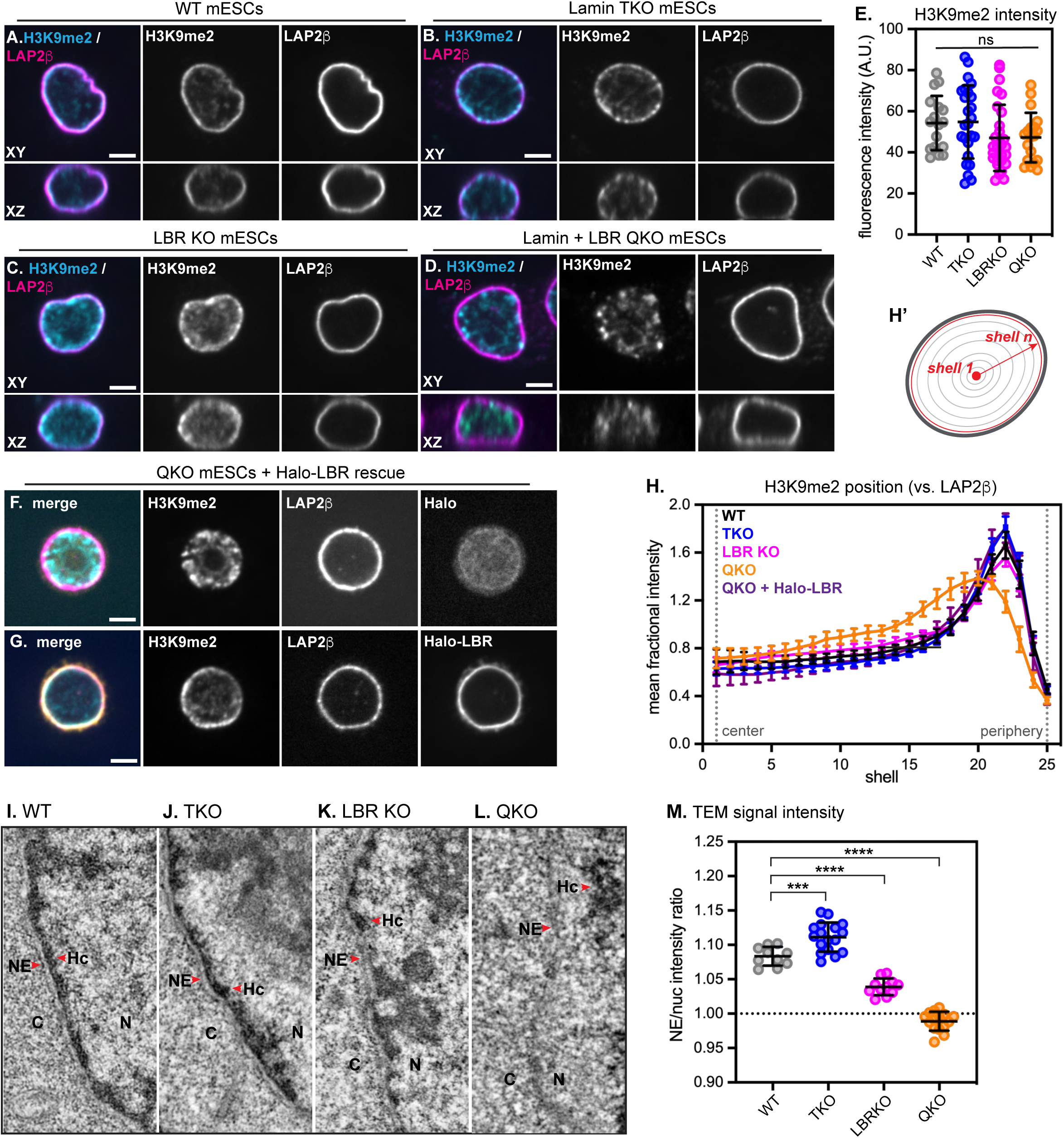
The lamins and LBR localize H3K9me2-marked chromatin to the nuclear periphery in mESCs. Immunofluorescence of H3K9me2 localization compared to the INM protein LAP2β in WT (A), lamin TKO (B), LBR KO (C), and lamin + LBR QKO (D) mESCs. A central z-slice (XY) and a central Y-slice (XZ) are shown. Scale bar, 5 μm. (E) H3K9me2 total fluorescence intensity per nucleus in WT (n = 17), TKO (n = 25), LBR KO (n = 28), and QKO mESCs (n = 19), p > 0.05 by one-way ANOVA. Bars indicate mean and standard deviation. (F) Immunofluorescence of H3K9me2 localization compared to the INM protein (LAP2β) in lamin + LBR QKO mESCS expressing Halo-NLS (F) or Halo-LBR (G) for 48 hours. (H,H’) Radial intensity analysis of H3K9me2 position in WT, TKO, LBR KO, and QKO mESCs (n = 25 nuclei per condition) and in QKO mESCs expressing Halo-LBR (n = 16 nuclei). **** p < 0.0001, WT vs QKO, shells 9-19, 22-24; * p < 0.05, WT vs TKO, shells 21-22; * p < 0.05, WT vs LBR KO, shells 14, 15, 16. Unpaired t-test used for all comparisons. Points indicate mean and error bars indicate 95% confidence intervals. See Figure S3 for full Halo-LBR rescue analysis and statistics. (I-L) Transmission electron microscopy showing 1.3 μm by 2.6 μm section of the nuclear periphery in WT (I), lamin TKO (J), LBR KO (K), and lamin + LBR QKO (L). C, cytoplasm; N, nucleus; NE, nuclear envelope; Hc, heterochromatin. (M) Quantification of relative TEM signal intensity ratio at the NE versus nucleoplasm (nuc), for WT (n = 10), LBRKO (n = 11), TKO (n = 17) and QKO (n = 15) mESCs. **** indicates p_adj_ < 0.0001 for WT vs LBRKO and WT vs QKO; *** indicates p_adj_ = 0.0002 for WT vs TKO by one-way ANOVA followed by Dunnett’s multiple comparisons test. Bars indicate mean and standard deviation.

We generated lamin + LBR quadruple knockout (QKO) and LBR knockout (LBR KO) mESCs by introducing frameshift indels into both alleles of the *Lbr* locus in lamin TKO mESCs and in WT littermate control mESCs, respectively (Fig. S1A, S1C-F). LBR KO, lamin TKO, and lamin + LBR QKO mESCs are each viable, pluripotent, and exhibit normal mESC colony morphology, albeit with a modest decrease in proliferation rate in QKO mESCs (Figure S1G-I).

We evaluated the effect of ablating the lamin and LBR proteins on the spatial organization of heterochromatin by quantifying the radial distribution of H3K9me2 (Fig. 1A-E, 1H) and compacted DNA (Fig. S2A-E), as well as by transmission electron microscopy (TEM) (Fig. 1I-M). These analyses revealed that H3K9me2-marked chromatin is displaced from the nuclear periphery and coalesces into nucleoplasmic foci in QKO mESCs (Fig. 1D, 1H), but not in lamin TKO (Fig. 1B, 1H) or LBR KO mESCs (Fig. 1C, 1H). Regions of more compacted chromatin can be visualized as regions of high relative intensity of DNA stain within the nucleus; while brightly stained, compacted chromatin is detectable at the nuclear periphery in WT, LBR KO, and TKO mESCs, compacted chromatin shifts away from the nuclear periphery and into the nucleoplasm in QKO mESCs (Fig. S2D,E). Further, while an electron-dense layer of heterochromatin is readily visible underneath the nuclear envelope in WT (Fig. 1I, 1M), lamin TKO (Fig. 1J, 1M), and LBR KO (Fig. 1K, 1M) mESCs, heterochromatin is undetectable underneath the NE in QKO mESCs (Fig. 1L-M) and is instead more prominent in the nucleoplasm (Fig. S2I). These data indicate that the lamins and LBR together control the spatial position of heterochromatin, consistent with previous work^30^; further, we determine that the organization of H3K9me2-modified chromatin depends on these proteins.

To understand the timescale of H3K9me2-marked chromatin displacement from the nuclear periphery in the absence of the lamins and LBR, we used a tetracycline-inducible RNAi system^37^ to induce the depletion of LBR to undetectable levels within 48 hours in TKO mESCs (Fig. S3F). Disorganization of H3K9me2-marked chromatin also became apparent within 48 hours (Fig. S3G-J). Re-establishment of H3K9me2 positioning could be rescued within 48 hours by expressing exogenous Halo-tagged LBR in QKO mESCs (Fig. 1G,H), but not by expressing the nucleoplasmic domain or transmembrane domain of LBR alone (Fig. S3C-E), which indicates that displacement of H3K9me2-marked chromatin from the nuclear periphery of lamin-depleted cells depends on LBR and is reversible.

### The genomic distribution of H3K9me2 is preserved in the absence of the lamins and LBR

We noted that ablating the lamins and LBR appears to affect the spatial distribution, but not the total abundance, of H3K9me2 (Fig. 1E). This observation suggests that deposition of H3K9me2 onto genomic loci occurs independently of its enrichment at the nuclear periphery. Since QKO mESCs have internalized their H3K9me2-marked chromatin, we asked whether this change in chromatin positioning affects the genomic position and abundance of H3K9me2. We used a monoclonal antibody with validated selectivity for this mark in genome-binding assays^14,15^ to perform CUT & RUN (<u>c</u>leavage <u>u</u>nder targets and release <u>u</u>sing <u>n</u>uclease) with spike-in control^38^. We then applied a four-state hidden Markov model (HMM)^39^, which identified domains lacking H3K9me2, domains with intermediate (class 1) H3K9me2 density, and domains with high (class 2) H3K9me2 density, as well as excluded blacklisted regions (Fig. 2A; Fig. S4; Table S1; see Methods). Altogether, we identified H3K9me2 domain-resident genes in WT mESCs that are closely concordant with previously published analyses of H3K9me2 (74% overlap; Fig. S4F-H) and LAD-resident genes (determined by LB1 ChIP-seq; 70% overlap; Fig. S4I-K) in mESCs in 2i + LIF culture conditions^14^. H3K9me2 domains appear similar across the genomes of WT, lamin TKO, LBR KO, and QKO mESCs (Fig. 2A), with similar numbers of class 1 and class 2 domains (Fig. 2B) that cover approximately 60% of the genome in kilobase-to megabase-long tracts (Figure 2C; Fig. S4B-C; ∼59%, ∼65%, ∼61%, and ∼57% of the genome covered in WT, LBR KO, TKO, and QKO mESCs, respectively). Approximately 66% of H3K9me2 domains are shared across all four genotypes (Fig. 2D), and over 70% are present in both WT and QKO mESCs (Fig. S4L). We noted modestly higher density of H3K9me2 within both class 1 and class 2 domains in QKO mESCs compared to other genotypes (Fig. 2E-F). Taken together, these data indicate that ablating the lamins and LBR disrupts the spatial positioning of H3K9me2 within the nucleus (Figure 1) but does not affect its deposition on the genome (Figure 2).

**Figure 2.**
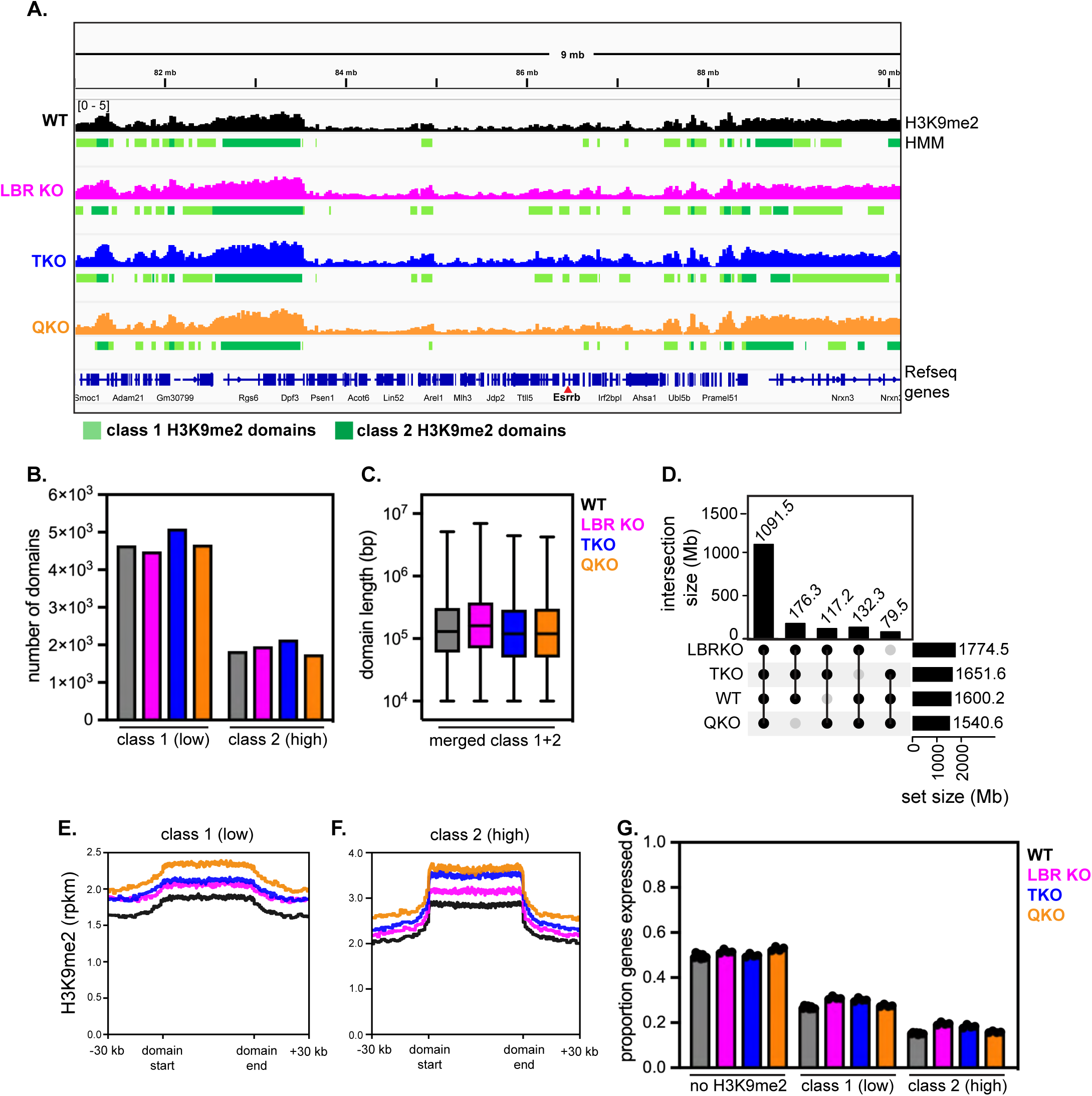
Genomic distribution of H3K9me2 is preserved after removal of the lamins and LBR. (A) Representative genome tracks and domain calls for H3K9me2 in WT, LBR KO, TKO, and QKO ESCs on a 9 Mb section of chromosome 12, including the *Esrrb* gene. Y axis range indicated at top left is the same for all tracks shown. Low K9me2 density “class 1” domains are marked in light green and high K9me2 density “class 2” domains are marked in dark green. (B) Number of HMM class 1 (low K9me2 density) and class 2 (high K9me2 density) domains called across genotypes. (C) Size of contiguous HMM class 1 (low K9me2 density) and class 2 (high K9me2 density) domains called across genotypes. (D) UpSet plot showing intersections of H3K9me2 domains between genotypes; the majority of domains are shared across all 4 genotypes, and a minority of domains are uniquely found in other 3-member groups of genotypes. Averaged density of H3K9me2 in all class 1 (E) and all class 2 (F) domains across genotypes. (G) Proportion of genes expressed (minimum of 5 TPMs) outside of H3K9me2 domains, within H3K9me2 class 1 domains, or within H3K9me2 class 2 domains across genotypes.

### Spatial displacement dysregulates H3K9me2-marked genes

Association of genes with the nuclear periphery is correlated with repression of transcription^8,9,11,40,41^. Consistently, we find a strong correlation between H3K9me2 modification, LAD residence, and low transcription (Fig. S4F-K, Fig. 2G); genes found within H3K9me2-marked domains are more likely to be repressed than genes found outside these regions, and the strength of repression correlates with the density of H3K9me2 (Fig. 2G). Transcriptional activation often correlates with displacement of genes from the nuclear periphery^12,14,42,43^, which suggests that positioning of genes at the nuclear periphery promotes repression. If this prediction is correct, heterochromatin displacement should weaken repression. Global transcriptional profiling by total RNAseq indicates that LBR ablation and lamin ablation each deregulate modest numbers of genes; in contrast, removal of both the lamins and LBR has a major synergistic effect on gene expression, with a bias toward upregulation (Fig. 3A; Table S2). We focused on changes in gene expression between lamin TKO mESCs (where heterochromatin positioning is intact) versus lamin + LBR QKO mESCs (where heterochromatin positioning is disrupted) (Fig. 3B); here, nearly twice as many genes are upregulated as are downregulated (983 genes > 2-fold up, 501 genes > 2-fold down). This outcome indicates that recruitment of heterochromatin to the nuclear periphery bolsters repression.

**Figure 3.**
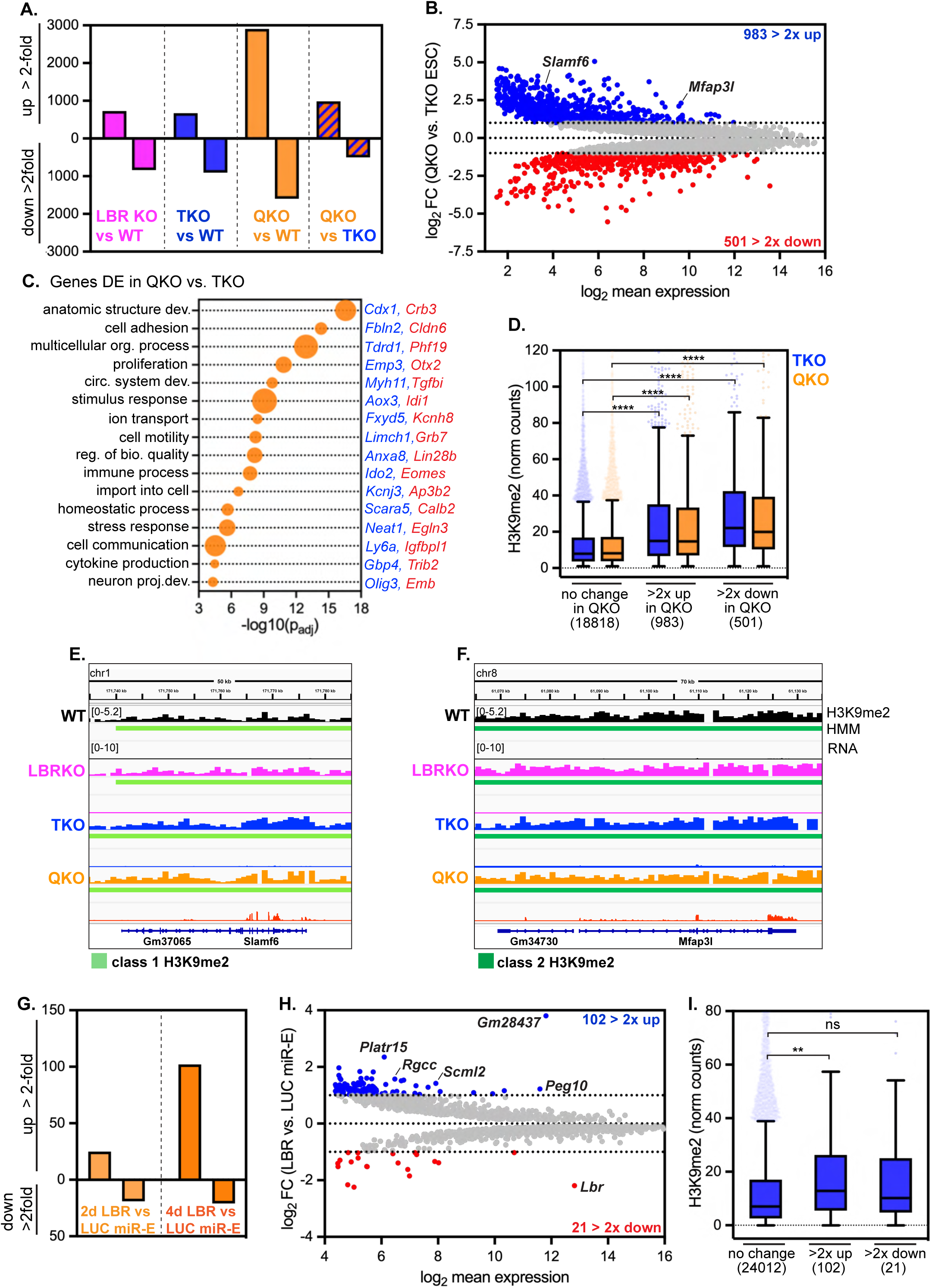
Removal of the lamins and LBR derepresses H3K9me2-marked and other genes. (A) Number of genes significantly differentially expressed at least 2-fold in all genotypes compared to wildtype mESCs, and in lamin + LBR QKO mESCs compared to lamin TKO mESCs. (B) MA plot comparing gene expression in QKO *versus* TKO mESCs. 5128 genes with a minimum *p_adj_* of 0.05 shown; 983 genes are upregulated at least 2-fold, while 501 genes are downregulated at least 2-fold. (C) Gene ontology analysis of all genes significantly differentially expressed by at least 2-fold in QKO vs. TKO mESCs. Biological process GO terms identified with gProfiler and reduced by ReVIGO. Selection of list shown as bubble plot where size corresponds to number of DE genes associated with that term (ranging from 60 to 459 genes). Representative upregulated and downregulated genes that intersect with each GO term are highlighted in blue and red, respectively. See also Supplementary Table 1 for full list of GO terms. (D) H3K9me2 levels (normalized counts) on genes with unchanged expression (*p_adj_* > 0.05, n=18818), genes upregulated at least 2-fold (n = 983) and genes downregulated at least 2-fold (n = 501) in QKO vs. TKO mESCs. **** indicates that comparisons indicated are significantly different (p < 0.0001) by Kruskal-Wallis multiple comparisons test with Dunn’s correction. Box (Tukey) plot center line indicates median; box limits indicate 25^th^ to 75^th^ percentiles; whiskers indicate 1.5x interquartile range; points indicate outlier values. (E) H3K9me2 and RNA levels of class 1 domain-resident gene *Slamf6* (E) and class 2 domain-resident gene *Mfap3l* (F) across genotypes. Y axis range indicated at the top left of each panel is the same for all tracks shown within panel. (G) Number of genes significantly differentially expressed at least 2-fold in lamin TKO mESCs expressing LBR versus LUC miR-E for 2 or 4 days. (H) MA plot comparing gene expression in lamin TKO mESCs expressing LBR or LUC miR-E for 4 days. 2055 genes with a minimum *p_adj_* of 0.05 shown; 102 genes are upregulated at least 2-fold, while 21 genes are downregulated at least 2-fold. (I) H3K9me2 levels (normalized counts) in TKO mESCs on genes with unchanged expression (*p_adj_* > 0.05, n=24012), genes upregulated at least 2-fold (n = 102) and genes downregulated at least 2-fold (n = 21) in TKO mESCs expressing LBR miR-E for 4 days. ** indicates significant difference (p = 0.0029) by Kruskal-Wallis multiple comparisons test with Dunn’s correction. Box (Tukey) plot center line indicates median; box limits indicate 25^th^ to 75^th^ percentiles; whiskers indicate 1.5x interquartile range; points indicate outlier values.

Genes involved in development, differentiation, signaling, and stress responses are dysregulated in QKO mESCs (Fig. 3C, Table S3), although expression of core pluripotency genes is maintained in all genotypes (Fig. S1I). H3K9me2 is enriched on dysregulated genes compared to unchanged genes (Fig. 3D), indicating that the transcriptional regulation of H3K9me2-marked genes is altered when heterochromatin is displaced. Further, inspection of derepressed loci in QKO mESCs reveals transcription within intact class 1 H3K9me2 domains (e.g. the class 1 resident upregulated gene *Slamf6,* Fig. 3E) and class 2 H3K9me2 domains (e.g. the class 2 resident upregulated gene *Mfap3l*, Fig. 3F). This de-repression occurs even though H3K9me2 domains are generally preserved in QKO mESCs (Fig. 2F) and are in fact accumulating even higher levels of H3K9me2 than in WT mESCs (Fig. 2D-E). These data indicate that some H3K9me2-marked genes are less effectively repressed when they are displaced from the nuclear periphery. In addition, some H3K9me2-marked genes are further repressed (Fig. 3D) while some non-H3K9me2-modified genes are also dysregulated(Fig. S5A), perhaps as secondary effects of heterochromatin disorganization.

To explore the initial consequences of heterochromatin displacement on gene expression, we analyzed the transcriptomes of lamin TKO mESCs after acute RNAi-mediated depletion of LBR for 2 or 4 days. These data revealed very few deregulated genes at day 2 (Fig. 3G; Fig. S5B; Table S4), when H3K9me2 disorganization is already apparent (Fig. S3); however, after 4 days of LBR depletion, 102 genes were upregulated more than 2-fold, while only 21 genes were downregulated more than 2-fold (one of which was *Lbr)* (Fig. 3G,H; Table S4), indicating a strong bias toward de-repression. These upregulated genes are significantly enriched for H3K9me2 compared to unaffected or downregulated genes (Fig. 3I). Altogether, we conclude that removal of the lamins and LBR causes the spatial displacement of H3K9me2-marked genes, followed by their de-repression.

### Heterochromatin displacement causes widespread de-repression of transposons

Deposition of H3K9me2 by G9a/GLP in mESCs^19,20^ promotes the effective repression of retrotransposons, including long interspersed elements (LINE1s) and endogenous retroviruses flanked by long terminal repeats (ERV LTRs)^21–25^. The role of heterochromatin positioning in repression of repeat elements is less clear, although LINE1s, ERV LTRs, and other transposable elements (TEs) reside within LADs^33,44^. As our data indicated that H3K9me2 is less able to repress a variety of genes when displaced from the nuclear periphery, we explored whether repression of TEs is also impaired. We applied TEtranscripts^45^ to detect and analyze the differential expression of multi-mapping RNAseq reads originating from ∼1200 distinct TE families that are integrated at numerous sites across the genome. LBR ablation had a minor effect on TE expression, with 7 TE families (0.5% of all) de-repressed at least 2-fold in LBR KO mESCs (Fig. 4A-B), while lamin ablation moderately induced TE expression, with 27 TE families (2% of all) de-repressed at least 2-fold in TKO mESCs (Fig. 4A,4C, Table S5). TEs that are de-repressed by lamin disruption alone include the ERV1 LTRs MMERGLN and MMERGLN-int and the L1MdA_I and L1MdA_II families of LINE1s (Fig. 4C). Co-depletion of the lamins and LBR dramatically elevated TE expression, with 338 TE families (27% overall) upregulated at least 2-fold in QKO mESCs compared to WT mESCs (Fig. 4A,4D, Table S5). TE classes including SINEs, LINEs, and satellites were affected in QKO mESCs, with LTR-containing retrotransposons the most widely de-repressed (38% of LTRs de-repressed; Fig. 4A). Within the LTR class, ERV-Ks including ERVB4_2-I and RLTR45-int are potently upregulated in QKO mESCs compared to both WT and TKO mESCs (Fig. 4C-D). Notably, ERV-K family members are similarly de-repressed when H3K9me2 is depleted via G9a/GLP inactivation in mESCs^24,25^. Depletion of the lamins and LBR appears to have a synergistic effect on LINE1 expression, as L1M3a, L1MdA_I, and L1MdA_II are further upregulated in QKO mESCs compared to TKO mESCs (Fig. 4C-D).

**Figure 4.**
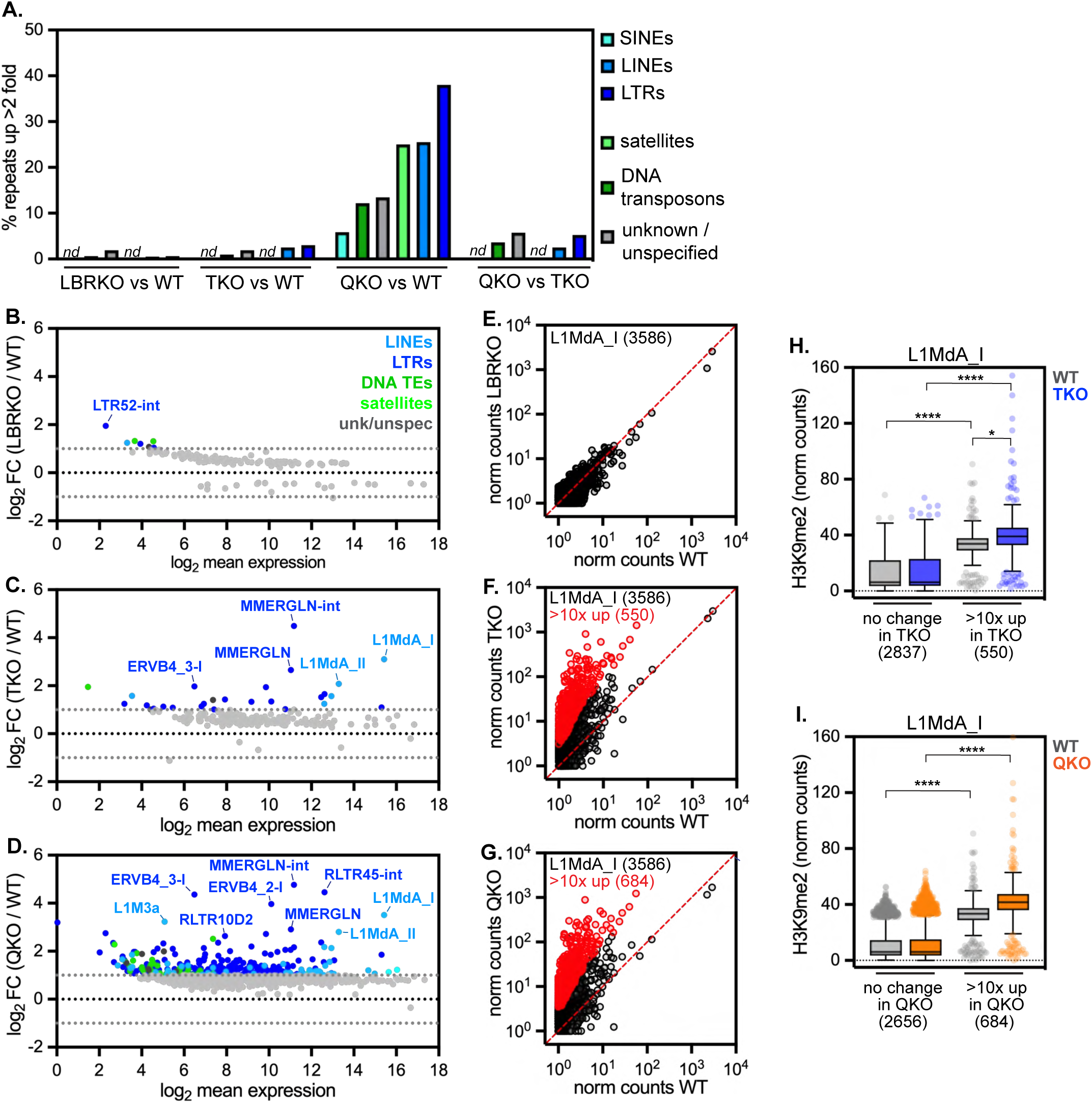
Ablation of the lamins and LBR allows pervasive transcription of transposons in spite of H3K9me2 modification. (A) Percentage of transposable element families de-repressed at least 2-fold in all genotypes compared to WT mESCs, determined by TEtranscripts. *nd* indicates not detected. MA plots comparing expression of ∼1200 TEs detected by TEtranscripts in (B) LBRKO versus WT mESCs, (B) TKO versus WT mESCs, and (D) QKO versus WT mESCs. All repeats without a significant change in between genotypes are gray; significantly differentially expressed TEs (minimum 2-fold change) are colored correspondingly to TE family. Normalized counts for 3586 uniquely mapped L1MdA_I LINE element genomic copies in (E) LBRKO versus WT mESCs, (F) TKO versus WT mESCs, and (G) QKO versus WT mESCs (plotted as log_10_(average + 1). L1MdA_I copies with >10-fold change and significant difference in expression (*p_adj_* < 0.05) in are colored in red. (H-I) Normalized counts from uniquely mapped reads of H3K9me2 on L1MdA_I LINE elements with unchanged expression versus those upregulated >10-fold in TKO mESCs (H) and in QKO mESCs (I). **** indicates p < 0.0001 and * indicates p = 0.0105 by one-way Kruskal-Wallis multiple comparisons test with Dunn’s correction. Box (Tukey) plot center line indicates median; box limits indicate 25^th^ to 75^th^ percentiles; whiskers indicate 1.5x interquartile range; points indicate outlier values.

Each TE family is present in numerous identical copies across the genome, but the expression of individual copies can be evaluated by searching for uniquely mapped transcripts that contain flanking genomic sequence. We used this approach to evaluate the expression of the large L1MdA_I LINE1 family^46^, which is present in 3586 copies in the mouse genome. While L1MdA_I copies remain lowly transcribed in WT and LBR KO mESCs (Fig. 4E), 550 individual L1MdA copies are upregulated at least 10-fold in TKO mESCs (Fig. 4F), while 684 copies are upregulated at least 10-fold in QKO mESCs (Fig. 4G). These data indicate that a significant proportion of the L1MdA_I LINE1 family is derepressed by depleting the lamins and LBR in mESCs.

To determine how quickly TEs become de-repressed when heterochromatin is displaced, we analyzed TE expression after acute RNAi-mediated depletion of LBR in lamin TKO mESCs for 2 or 4 days. This comparison is useful in spite of the fact that some TEs are de-repressed in lamin TKO mESCs, as an additional 52 TE families are derepressed in lamin + LBR QKO mESCs compared to lamin TKOs (Fig. 4A). Strikingly, 42 uniquely mapped TE copies were potently derepressed greater than 5-fold within 4 days of LBR depletion in TKO mESCs (Fig. S5C-D), indicating that TE de-repression follows H3K9me2 displacement.

The effects of heterochromatin displacement on TE expression are reminiscent of the effects of deleting H3K9 methyltransferases, which reduces H3K9me2/3 on TEs and allows their transcription^23–25,47,48^. However, our analyses indicate that H3K9me2 is generally maintained on the same genomic loci when displaced from the nuclear periphery (Fig. 2). To evaluate H3K9me2 on TEs in the absence *versus* presence of the lamins and LBR, we analyzed uniquely mapped H3K9me2 CUT & RUN reads that included L1MdA_I LINE1 sequences. This analysis indicated that (i) L1MdA_I copies that are sensitive to the loss of the lamins and LBR are significantly more highly modified by H3K9me2 than unaffected L1MdA_I copies (Fig. 4H, 4I); and (ii) that these same L1MdA_I copies remain densely decorated by H3K9me2 in TKO and QKO mESCs and in fact gain even more H3K9me2 in mutant mESCs compared to WT mESCs (Fig. 4H, 4I). Therefore, similarly to our analyses of genes (Fig. 3), heterochromatin displacement allows the transcription of TEs in spite of their persistent H3K9me2 modification.

### Heterochromatin positioning enables the transition from naïve to primed pluripotency

Our data indicate that H3K9me2 loses the capacity to repress the transcription of protein-coding genes (Fig. 3) and of TEs (Fig. 4) when displaced from the nuclear periphery. These observations lead us to hypothesize that the recruitment of H3K9me2-marked chromatin to the nuclear periphery strengthens H3K9me2-mediated repression. To further test the influence of heterochromatin positioning on the function of H3K9me2, we asked whether it is required for the transition from naïve to primed pluripotency, a developmental transition that can only be completed if G9a and GLP are upregulated to expand H3K9me2 across the genome^24,49^. *In vivo*, this transition occurs when the inner cell mass of an embryo exits naïve pluripotency and enters the epiblast state. While naïve pluripotency is approximated in culture by mESC growth in 2i + LIF conditions, induction of epiblast-like cells (EpiLCs) can be achieved by treatment of mESCs with FGF and Activin A growth factors for 24-48 hours^50,51^ (Fig. 5A). LBR KO and lamin TKO mESCs each exhibited a moderate defect in growth and/or survival during the transition to EpiLCs, lamin + LBR QKO mESCs were severely impaired in their ability to differentiate (Fig. 5B,5C), and only ∼20% completed this transition. Further, EpiLC loss is an immediate consequence of lamin + LBR co-depletion, as acute depletion of LBR by RNAi in lamin TKOs impaired EpiLC differentiation (Fig. 5D).

**Figure 5.**
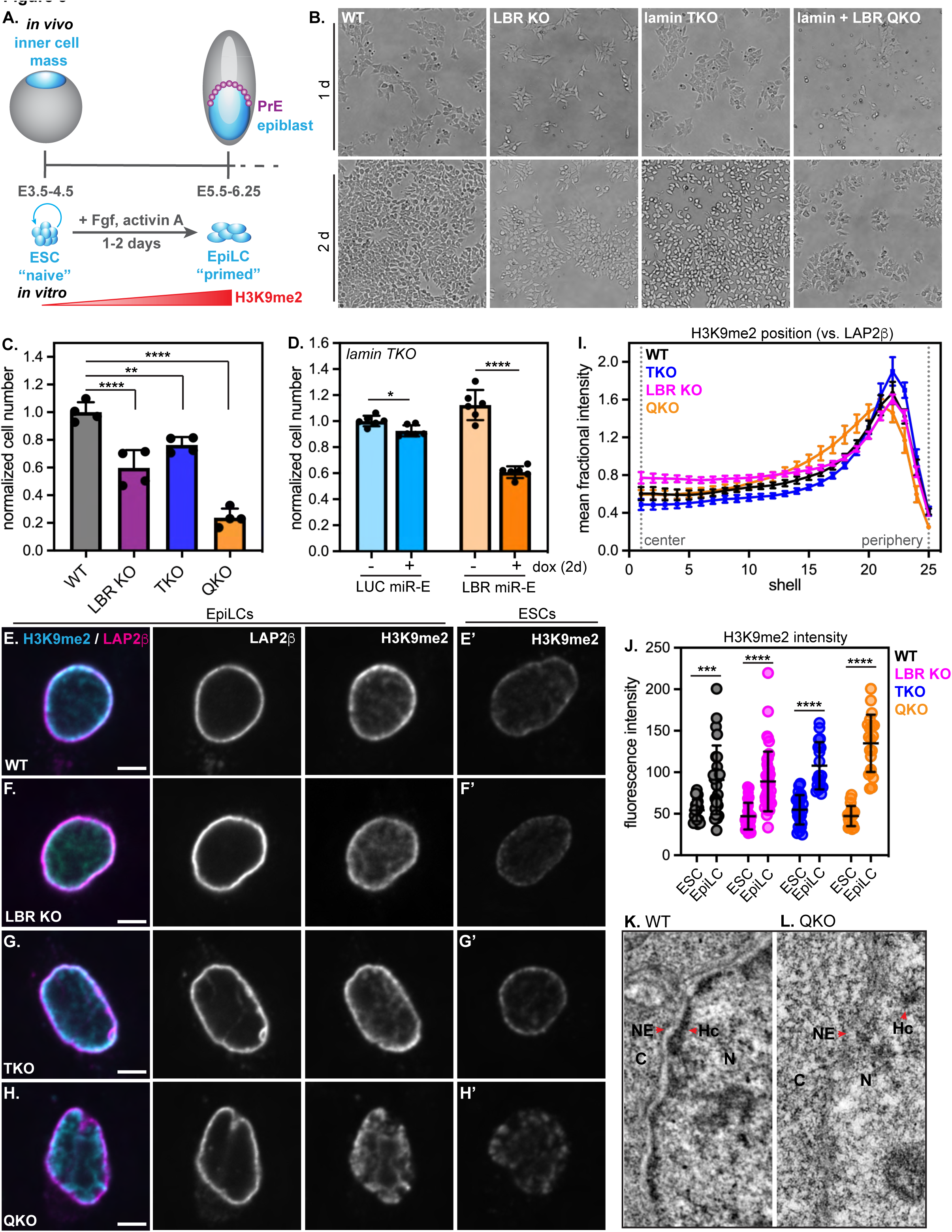
Heterochromatin positioning by the lamins and LBR is essential for EpiLC survival. (A) Diagram of naïve (ICM / mESC) to primed (epiblast / EpiLC) developmental transition *in vivo* and its approximation *in vitro.* (B) Representative brightfield microscopy images of WT, LBR KO, lamin TKO, and lamin + LBR QKO mESCs after 1 day (top panel) or 2 days (bottom panel) of culture in EpiLC differentiation conditions. (C) Normalized cell numbers after 2 days of culture in EpiLC differentiation conditions (normalized to WT; n = 4 replicates per genotype); columns indicate mean, error bars indicate standard deviation. **** indicates p_adj_ < 0.0001 for WT vs LBRKO and WT vs QKO; ** indicates p_adj_ = 0.0058 for WT vs TKO by one-way ANOVA followed by Dunnett’s multiple comparisons test. (D) Normalized cell numbers of lamin TKO mESCs after 2 days of culture in EpiLC differentiation with (+) or without (-) co-induction of LUC or LBR miR-E. Columns indicate mean and error bars indicate standard deviation. n = 6 replicates from 2 independent clones shown. *, p < 0.05 and ****, p < 0.0001 by unpaired t-test. Immunofluorescence of H3K9me2 localization compared to the INM protein LAP2β in WT (E-E’), LBR KO (F-F’), lamin TKO (G-G’), and lamin + LBR QKO (H-H’) cells. (E-H) show EpiLC samples while (E’-H’) show mESC samples stained and imaged in parallel. Central z-slices (XY) shown. Scale bar, 5 μm. (I) Radial intensity analysis of H3K9me2 position *versus* LAP2b in WT, TKO, LBR KO, and QKO EpiLCs. **** p < 0.0001, WT vs QKO, shells 14-19, 24-25. (J) H3K9me2 total fluorescence intensity per nucleus in WT (n = 26), TKO (n = 18), LBR KO (n = 41), and QKO (n = 20) EpiLCs. *** p < 0.001, **** p < 0.0001, unpaired t-test. Transmission electron microscopy showing 1.3 μm by 2.6 μm section of the nuclear periphery in WT (K) and lamin + LBR QKO (L) EpiLCs. C, cytoplasm; N, nucleus; NE, nuclear envelope; Hc, heterochromatin.

In addition to their shared role in heterochromatin positioning, LBR and the lamins perform other important functions that could potentially be required to sustain EpiLC survival. We tested the relevance of these other functions to EpiLC survival as follows. Because LBR participates in cholesterol biosynthesis^52^, we evaluated cholesterol levels in WT and mutant mESCs and EpiLCs. Cholesterol levels are unaffected by LBR ablation (Fig. S6A), likely because of the redundant activity of the TM7SF2 enzyme^52^. The effect of deleting LBR on EpiLC survival is thus not a consequence of cholesterol depletion. mESCs flatten, migrate, and proliferate as they transform into EpiLCs (Fig. 5B). Because the lamins and heterochromatin each absorb forces on the nucleus^53^, it is possible that mutant nuclei might not be able to endure the forces that arise from cellular flattening during differentiation. To test whether cellular flattening impairs survival of lamin + LBR QKO mESCs, we compared the viability of mESCs growing on a low-attachment substrate (gelatin), where cells form rounded colonies, to a higher-attachment substrate (Cultrex), where cells form more flattened and adherent colonies (Fig. S6B-C). We found no difference in cell viability between these conditions, indicating that cellular flattening itself does not impair survival of QKO mESCs.

We re-evaluated the spatial positioning of heterochromatin within EpiLC nuclei and found that similarly to QKO mESCs (Fig. 1), heterochromatin displacement from the nuclear periphery of QKO EpiLCs is apparent by H3K9me2 immunostaining (Fig. 5E-I), by Hoechst staining of DNA compaction (Fig, S7A-D, E), and by TEM (Fig. 5K-L; Fig. S7F-J). Altogether, these data confirm that the lamins and LBR are also required for peripheral heterochromatin positioning in EpiLCs and demonstrate that H3K9me2 positioning is required for EpiLC viability.

### Heterochromatin positioning influences remodeling of H3K9me2 in primed pluripotency

While our microscopy analyses indicate that H3K9me2 is displaced from the nuclear periphery in QKO EpiLCs (Fig. 5H,I), we noted that H3K9me2 abundance appears to increase during the transition from naïve to primed pluripotency in all genotypes (Fig. 5E-H’; Fig. 5J). To quantify H3K9me2 abundance across the genome in EpiLCs, we again applied spike-in-controlled H3K9me2 CUT & RUN and identified domains of absent, intermediate (class 1), or high (class 2) H3K9me2 intensity with a 4-state HMM (Fig. 6). This spike-in-controlled analysis demonstrated expansion of H3K9me2 across the genome in WT EpiLCs *versus* mESCs that was apparent by visual inspection (Fig. 6A), by quantifying total genome coverage (Fig. S9C vs Fig. S4B; ∼66% of genome within H3K9me2 domains in EpiLCs vs. ∼59% in ESCs), by quantifying the median contiguous length of H3K9me2 domains in WT ESCs vs. EpiLCs (Fig. 6B; 130 kb in WT ESCs vs 190 kb in WT EpiLCs), and by tracking the net flow of genes into H3K9me2 domains in WT EpiLCs *versus* ESCs (Fig. 6C; 18729 genes move into H3K9me2 domains in EpiLCs; Table S6). Interestingly, the establishment of EpiLC-specific H3K9me2 domains is not accompanied by widespread repression of these newly modified genes; instead, most EpiLC H3K9me2 genes are unchanged in expression, while small numbers of genes are upregulated or downregulated (Fig. 6D). While a lower proportion of genes within constitutive H3K9me2 domains (those shared between mESCs and EpiLCs) are transcribed, a high proportion of genes within EpiLC H3K9me2 domains are transcribed (Fig. 6E). These observations suggest that H3K9me2 is overall less repressive and performs a distinct function in EpiLCs compared to mESCs. Altogether, these data indicate that H3K9me2 rapidly expands during the transition into primed pluripotency without immediately inducing widespread repression of newly modified loci.

**Figure 6.**
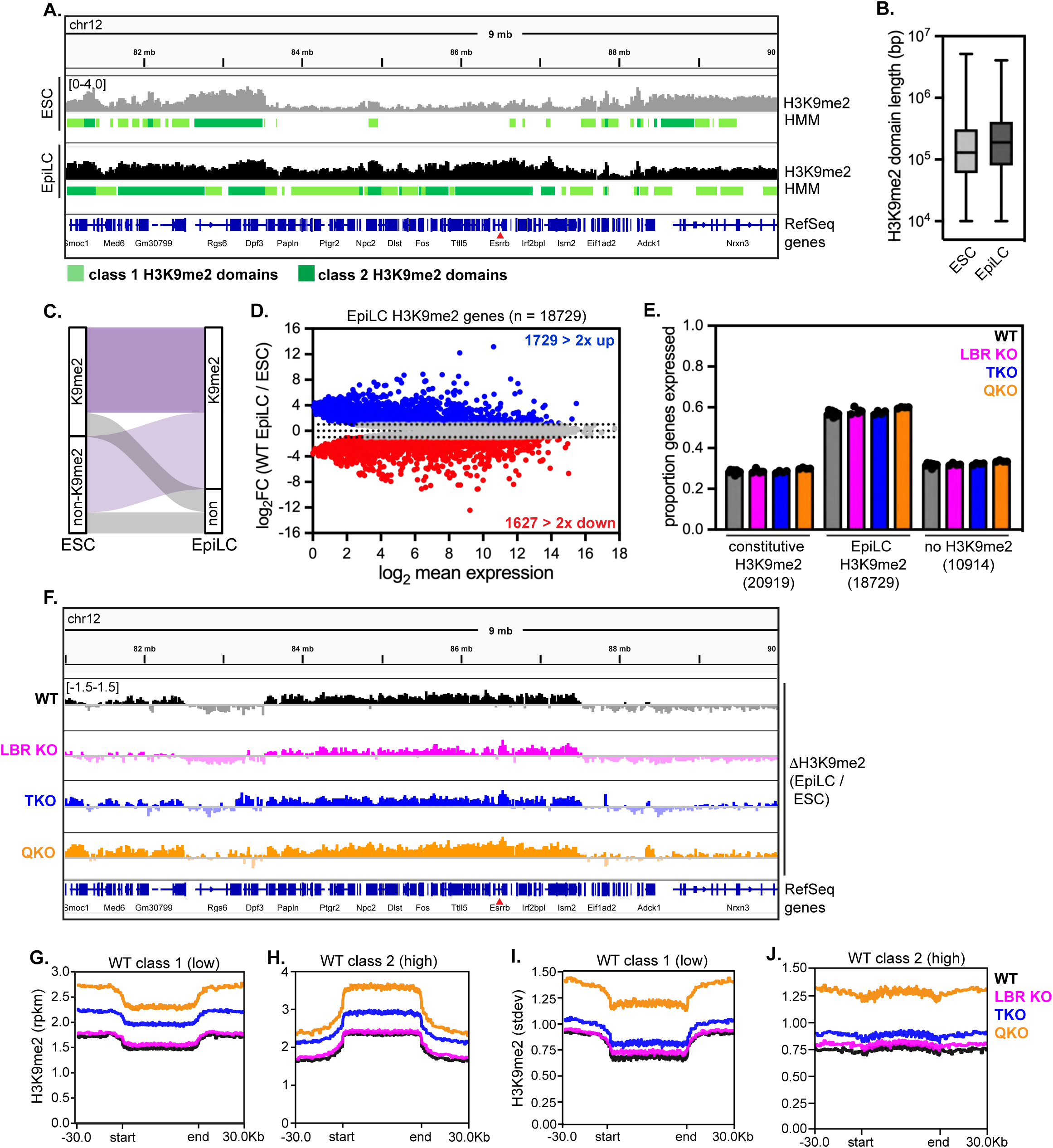
Abnormal deposition of H3K9me2 in Lamin + LBR KO EpiLCs. (A) Representative genome tracks and domain calls for H3K9me2 in WT ESCs and EpiLCs on a 9 Mb section of chromosome 12, including the *Esrrb* gene. Y-axis range indicated at top left is the same for all tracks shown. Low K9me2 density “class 1” domains are marked in light green and high K9me2 density “class 2” domains are marked in dark green. (B) Median contiguous size of HMM class 1 (low K9me2 density) and class 2 (high K9me2 density) domains in WT ESCs versus EpiLCs. (C) Alluvial plots showing movement of genes into and out of H3K9me2 domains as WT ESCs differentiate into EpiLCs. Genes found in H3K9me2 domains in both ESCs and EpiLCs are referred to as “constitutive” (dark purple, 20919 genes)) while genes that move into H3K9me2 domains in EpiLCs are referred to as “EpiLC H3K9me2” (light purple, 18729 genes). 10914 genes are not included in H3K9me2 domains in EpiLCs. (D) MA plot comparing expression of EpiLC H3K9me2 genes in WT EpiLCs versus mESCs. Of 18729 genes that move into EpiLC H3K9me2 domains, 7590 with *p_adj_* < 0.05 are plotted; of these, 1729 genes are upregulated at least 2-fold and 1627 are downregulated at least 2-fold. (E) Proportion of genes expressed (minimum of 5 TPMs) within constitutive H3K9me2 domains, within EpiLC H3K9me2 domains, or outside of H3K9me2 domains across genotypes. (F) Difference maps of H3K9me2 signal in EpiLCs vs. ESCs across a 9 Mb section of chromosome 12. Y-axis range indicated at top left is the same for all tracks shown. (G-H) Averaged density of H3K9me2 in class 1 (G) and class 2 (H) domains (domain coordinates identified in WT EpiLCs). (I-J) Standard deviation of H3K9me2 signal in class 1 (I) and class 2 (J) domains (domain coordinates identified in WT EpiLCs).

We next applied spike-in-controlled CUT & RUN to mutant EpiLCs. Similarly to WT EpiLCs, expansion of H3K9me2 across the genome was readily apparent in LBR KO, lamin TKO, and lamin + LBR QKO EpiLCs (Fig. 6F). Our domain calling approach indicated that in WT, LBR KO and lamin TKO EpiLCs, H3K9me2 domains increase in median length and incorporate new genes (Fig. 6B-C; Fig. S9G-I) that remain transcriptionally active (Fig. 6E). However, domain calling identified short and fragmented H3K9me2 domains in QKO EpiLCs, even across genomic regions with high overall H3K9me2 signal (Fig. S9C,E,F). To avoid the confounding effects of variable HMM performance across samples, we compared H3K9me2 density within H3K9me2 domains identified in WT EpiLCs. Within these domain borders, QKO EpiLCs accumulate H3K9me2 to significantly higher levels than do other genotypes (Fig. 6G,H). In addition, H3K9me2 signal variance is significantly greater both genome-wide and within WT H3K9me2 domain borders in QKO EpiLCs compared to other genotypes (Fig. S9B; Fig. 6I,J). We surmise that the abnormally variable pattern of H3K9me2 deposition across the genome in QKO EpiLCs interferes with HMM domain calling. Taken together, these analyses indicate that heterochromatin displacement alters both the level and the pattern of H3K9me2 deposition along the genome during the transition from naïve to primed pluripotency.

### Heterochromatin positioning silences transposons and alternative cell fate genes in EpiLCs

We next evaluated the consequences of H3K9me2 displacement on the dynamic transcriptional landscape of EpiLCs. QKO EpiLCs exhibit widespread de-repression of TEs, with 298 TEs (24% overall) upregulated at least 2-fold compared to WT EpiLCs, and 224 TEs (18% overall) upregulated at least 2-fold compared to TKO EpiLCs (Fig. S10A, Table S7). In contrast, LBR KO and lamin TKO EpiLCs maintain repression of most TEs (Fig. 7A; Fig. S10A; Table S7). LINEs, satellites, and ERV LTRs are broadly de-repressed in QKO EpiLCs (Fig. 7A; Fig. S10D).

**Figure 7.**
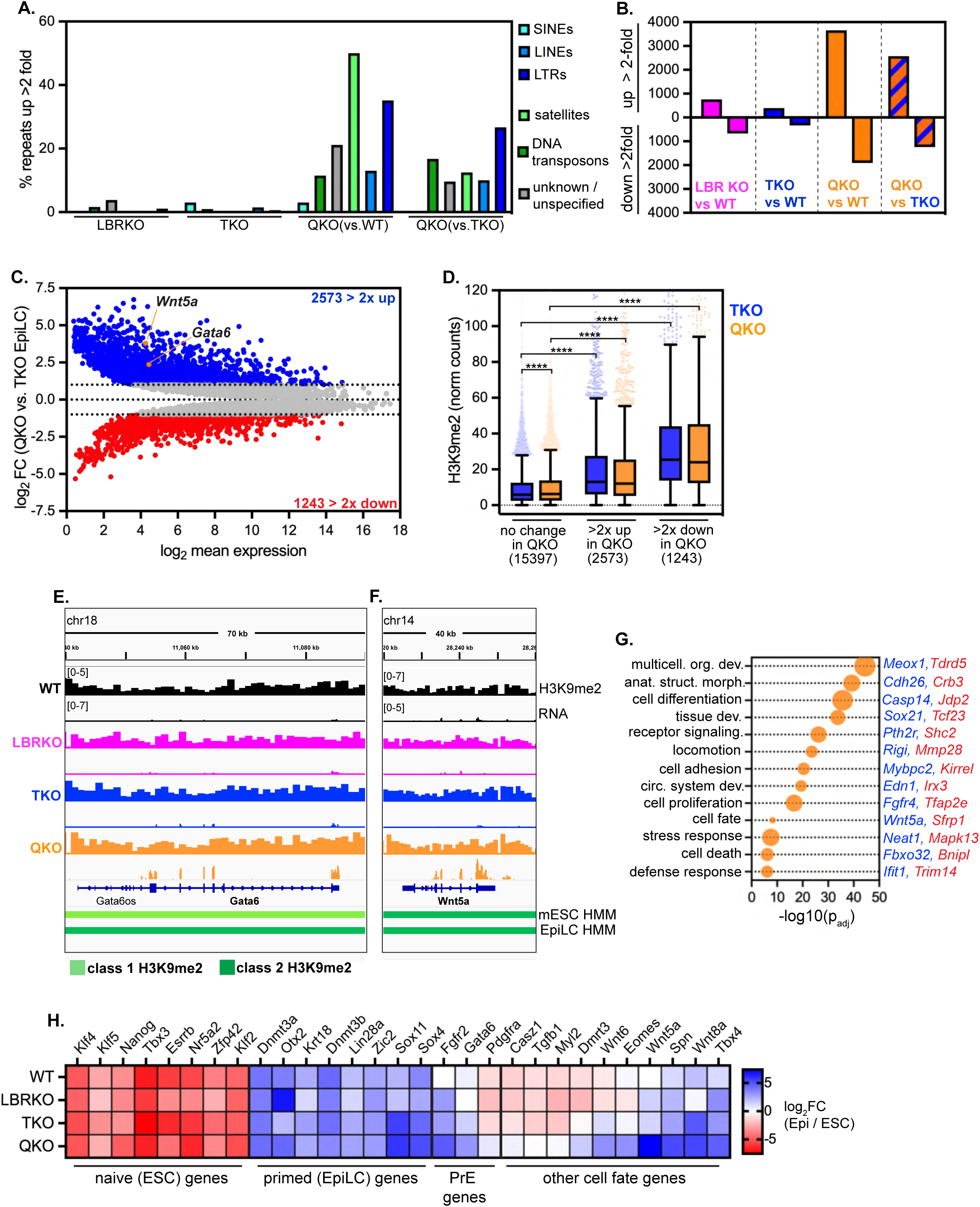
Removal of the lamins and LBR derepresses genes and transposons and impairs lineage restriction in EpiLCs. (A) Percentage of TE families de-repressed at least 2-fold in LBR KO, lamin TKO, and lamin + LBR QKO EpiLCs *versus* WT EpiLCs, and in lamin + LBR QKO EpilCs *versus* TKO EpiLCs. (B) Number of genes significantly differentially expressed at least 2-fold in all genotypes compared to wildtype EpiLCs, and in lamin + LBR QKO EpiLCs compared to lamin TKO EpiLCs. (C) MA plot comparing gene expression in QKO *versus* TKO EpiLCs. 10051 genes with a minimum *p_adj_* of 0.05 shown; 2573 genes are upregulated at least 2-fold, while 1243 genes are downregulated at least 2-fold. Representative H3K9me2-modified and de-repressed genes *Wnt5a* and *Gata6a* highlighted. (D) Analysis of H3K9me2 modification levels (normalized counts) on genes unchanged in expression between TKO and QKO EpiLCs (n = 15397), genes upregulated at least 2-fold in QKO EpiLCs (n = 2573), and genes downregulated at least 2-fold in QKO EpiLCs (n = 1243). **** indicates that comparisons indicated are significantly different (p < 0.0001) by Kruskal-Wallis multiple comparisons test with Dunn’s correction. Box (Tukey) plot center line indicates median; box limits indicate 25^th^ to 75^th^ percentiles; whiskers indicate 1.5x interquartile range; points indicate outlier values. (E-F) H3K9me2 and RNA levels of H3K9me2 domain genes *Gata6* (E) and *Wnt5a* (F) across genotypes. Y axis ranges indicated at the top left of each panel are the same for all lower tracks within panel. (G) Gene ontology analysis of all genes significantly differentially expressed by at least 2-fold in QKO vs. TKO EpiLCs. Biological process GO terms identified with gProfiler and reduced by ReVIGO. Selection of list shown as bubble plot where size corresponds to number of DE genes associated with that term (ranging from 64 to 697 genes). Representative upregulated and downregulated genes that intersect with each GO term are highlighted in blue and red, respectively. See also Supplementary Table 2 for full list of GO terms. (H) Heatmap showing expression of selected genes associated with naïve pluripotency, primed pluripotency, primitive endoderm, or other cell fates in each genotype. Heatmap values indicate log2(fold change) in expression between EpiLCs and ESCs.

To further explore the extent of TE de-repression, we analyzed unique transcripts originating from the L1MdA_I LINE1 family; while this family remains repressed in LBR KO EpiLCs (Fig. S10E), 96 copies were upregulated at least 10-fold in lamin TKO EpiLCs (Fig. S10F), while 229 copies were upregulated at least 10-fold in QKO EpiLCs compared to WT EpiLCs (Fig. S10G). Overall, L1MdA_I expression was more moderate in EpiLCs compared to mESCs (Fig. 4), suggesting that additional TE repressive mechanisms may be induced in primed pluripotency, or that only EpiLCs expressing moderate levels of TEs survive. Nevertheless, derepressed L1MdA_I copies are highly H3K9me2-modified in both wild type and mutant EpiLCs, while unaffected L1MdA_I copies are not (Fig. S10H-I), indicating that some TEs reside in constitutive H3K9me2 domains in both mESCs and EpiLCs, and that spatial displacement of H3K9me2 allows the selective de-repression of these TEs.

LBR KO and lamin TKO EpiLCs each exhibit modest changes to gene expression, while co-depletion of the lamins and LBR has a synergistic effect on gene expression (Fig. 7B; Table S8), similarly to our observations in naïve mESCs (Fig. 3). Lamin + LBR QKO EpiLCs exhibit a bias toward de-repression of genes compared to lamin TKO EpiLCs (Fig. 7B, 7C), with 2573 genes upregulated at least 2-fold and 1243 genes downregulated at least 2-fold. These dysregulated genes are enriched for the H3K9me2 modification compared to unaffected genes, indicating that disrupting heterochromatin recruitment affects the regulation of H3K9me2-modified genes (Fig. 7D). Inspection of transcription within H3K9me2 domains revealed examples of developmentally important genes that are de-repressed in QKO EpiLCs yet reside within H3K9me2 domains, such as the transcription factor *Gata6* (Fig. 7E) and the morphogen *Wnt5a* (Fig. 7F).

Genes dysregulated in QKO EpiLCs include components of immune response, host defense, stress, and apoptosis pathways (Fig. 7G; Table S9). Notably, genes involved in the innate immune response to viral RNA are upregulated, including *Ifit1* and *Rigi*; transcription of these genes is induced by TE activity^54,55^ and can drive apoptosis^56^. Dysregulated genes also participate in a range of cell fate determination, morphogenesis, and differentiation networks, suggesting that developmental progression is abnormal (Fig. 7G; Table S9). We investigated the regulation of cell fate in mutant EpiLCs more closely (Fig. 7H), which indicated that QKO EpiLCs succeed in silencing naïve pluripotency genes and activating primed/epiblast stage pluripotency genes. Therefore, QKO mESCs can undergo a cell state change in response to a pro-differentiation stimulus. However, we noted that in addition to appropriately inducing epiblast-stage genes, QKO EpiLCs abnormally express various alternative cell fate genes (Fig. 7H). These include markers of the primitive endoderm, such as *Fgfr2, Gata6,* and *Pdgfra*; this extra-embryonic lineage should be mutually exclusive with the epiblast fate^57,58^ (Fig. 7H). QKO EpiLCs also abnormally express a range of other lineage-specific morphogens and transcription factors, such as *Wnt5a, Wnt6, Wnt8a, Tgfb1,* and *Eomes* (Fig. 7H). This outcome suggests that while recruitment of heterochromatin to the nuclear periphery is not required for the transition from naïve to primed pluripotency *per se*, it enables the effective specification of a lineage-specific gene expression program. Altogether, we conclude that the lamins and LBR together control both the spatial positioning and the repressive capacity of H3K9me2 to shape cell fate decisions during early mammalian development.

## Discussion

### The lamins and LBR exert broad influence on heterochromatin organization and function

The parallel roles of lamin A/C and LBR in heterochromatin organization were first described over 10 years ago^30^. We demonstrate that the lamins and LBR together exert broad control on heterochromatin organization and function in pluripotent cells. Strikingly, we observe that the dense layer of compacted, electron-dense heterochromatin that is a hallmark of most eukaryotic cells disperses when the lamins and LBR are ablated (Fig. 1L). H3K9me2-marked loci are also displaced from the nuclear periphery and into nucleoplasmic foci in cells lacking these proteins (Fig. 1D, 1F). This phenotype bears some similarity to the gradual intranuclear coalescence of heterochromatin that occurs in several types of neurons that downregulate lamin A/C and LBR as they differentiate^28,30,59^. Neuronal chromatin inversion occurs without any disruption to heterochromatin-mediated repression^28,59^, while our data indicate that heterochromatin displacement in pluripotent cells causes dramatic transcriptional dysregulation. We surmise that the high proliferation rate of mESCs prevents the complete coalescence of heterochromatin into a single intranuclear focus, as occurs in neurons over several weeks after cell cycle exit^30^. Alternatively, additional chromatin changes such as relabeling of H3K9me2 loci with H3K9me3 and/or presence of distinct H3K9me2/3 binding proteins may promote heterochromatin coalescence in neurons^28^ but not in mESCs (Fig. 2). Nevertheless, our findings indicate that heterochromatin reorganization can be induced by the removal of the lamins and LBR in pluripotent cells. Further, our data establish for the first time that the unique enrichment of H3K9me2-marked chromatin underneath the nuclear periphery is maintained by the redundant function of the lamin and LBR proteins.

Our data indicate that the lamins can sustain chromatin organization in the absence of LBR and vice versa. While LBR is much more highly expressed than lamin A/C in pluripotent cells, low levels of lamin A/C have been recently shown to influence gene expression in naïve pluripotency^60^. LBR alone can drive peripheral heterochromatin positioning when ectopically expressed in various cell types, while ectopically expressed Lamin A/C cannot^30^; this implies that Lamin A/C’s heterochromatin tethering function is mediated by additional factors with variable expression levels across tissues, while LBR can either directly tether heterochromatin, or alternatively work through ubiquitously expressed intermediary protein(s). Various lamin-bound proteins such as LAP2β, HDAC3, PRR14, and others are candidates that could promote lamin-mediated heterochromatin tethering^9,14,61,62^, while LBR binds to the H3K9me2/3-binding protein HP1^63^ and can also interact with histone tails via its Tudor domain^64,65^. In the future, we will dissect how these interactions contribute to heterochromatin tethering in the sensitized background of lamin-null or LBR-null mESCs.

### The nuclear periphery influences the repressive capacity of H3K9me2

We show that disruption of the lamin and LBR heterochromatin tethering proteins weakens the repression of H3K9me2-modified loci even though their H3K9me2 modification is preserved (Fig. 3B,D). While H3K9me2 has a well-established capability to repress transcription^19,20^, our data demonstrate that nuclear spatial position influences the function of H3K9me2 and is consistent with recent indications that the repressive capacity of H3K9me2 is context-dependent. For instance, chromatin bearing H3K9me2 alone is relatively more permissive to transcription than H3K9me2-modified and lamina-associated regions of the genome^27^. The function of H3K9me2 also appears to change during the differentiation of photoreceptor neurons, where pre-existing repressive H3K9me2 is gradually converted to H3K9me3 to preserve repression, while euchromatic regions of the genome acquire H3K9me2 but remain transcribed^28^. Notably, this shift in H3K9me2’s function is accompanied by the downregulation of the lamin A/C and LBR heterochromatin tethers^30^; similarly, we demonstrate that removal of the lamins and LBR decreases the repressive capacity of H3K9me2. By using an inducible RNAi system to acutely remove LBR in lamin-null mESCs, we determined that detachment of H3K9me2 is followed by de-repression of H3K9me2-marked genes within days (Fig. 3G-I). Based on these observations, we propose that recruitment to the nuclear periphery favors repression of H3K9me2-marked loci, while displacement from the nuclear periphery enables expression of H3K9me2-marked loci.

Heterochromatin tethering could enhance repression by several potential mechanisms. Repression may be induced by steric occlusion if binding of H3K9me2 and oligomerization of tethering proteins together promote compaction of chromatin domains. Similarly, tethering to the nuclear periphery may limit turnover of nucleosomes to enable long-term memory of chromatin state, as has been demonstrated in *S. pombe*^7^. Finally, repression at the periphery may be achieved by the addition or removal of other chromatin marks to consolidate repression. For instance, tethering of a locus to the nuclear periphery is often accompanied by histone deacetylation^9,40^.

### The nuclear periphery maintains repression of transposons

TEs are enriched in LADs in both pluripotent and differentiated cells^33,44^, and our data indicate that lamin disruption allows the expression of some TE families, particularly in naive pluripotency (Fig. 4A, C). While altered gene expression was previously reported in lamin TKO mESCs, the expression of TEs in these cells was not evaluated^36^. Intriguingly, alterations to the lamina that occur in aged and senescent cells have been linked to displacement and/or activation of retrotransposons^66–69^, suggesting that the lamina may also repression of TEs in differentiated cells.

H3K9me2 and H3K9me3 each play a major role in repression of TEs^48^, and we find that displacement of H3K9me2 from the nuclear periphery by ablation of both the lamins and LBR leads to pervasive activation of both retrotransposons and DNA transposons. By analyzing individual genomic copies of the large L1MdA_I LINE1 family, we determined that H3K9me2-marked TEs are sensitive to activation upon H3K9me2 displacement, while TEs that lack H3K9me2 are not affected (Fig. 4H-I). Strikingly, the effect of H3K9me2 displacement on TE expression is comparable to the effect of preventing the deposition of H3K9me2. For instance, ablating the H3K9me2/3-depositing enzyme SETDB1 activates a subset of ERV LTRs^47^ while ablating the H3K9me2 enzyme G9a/GLP or ablating all H3K9me2 enzymes together(SETDB1, SETDB2, G9a, and GLP) de-represses many ERV LTRs as well as LINE1s^21–25,48^. We find that acute depletion of LBR in lamin TKO mESCs induces TE expression within 2-4 days (Fig. S5C-D), indicating that TE activation is a rapid response to heterochromatin displacement. Therefore, we conclude that peripheral positioning is required for the effective repression of H3K9me2-modified TEs in pluripotent cells.

TE activation can have wide-ranging effects on genome function and stability (reviewed in^70^). TEs can act as alternative enhancers, promoters, or exons to create novel lncRNAs or protein-coding genes^22,47,71^. Younger TE families that retain the ability to undergo transposition will be unleashed to do so if they become transcriptionally active and will perturb the loci they insert into. Several young and mobile TE families are derepressed when heterochromatin is displaced in QKO cells. These include the L1MdA_I LINE1 family, which is the youngest LINE1 family in the mouse genome^46^; and ERV LTRs, many of which undergo transposition in the mouse genome^72^. The mobilization of autonomous TEs (such as LINE1s) can in turn induce the activity and/or transposition of non-autonomous TEs such as SINEs^70^. Altogether, this cascade of effects initiated by TE activation may explain why we observe changes in both gene and TE expression that include but are not limited to H3K9me2-modified loci.

Mechanisms of TE repression change as cells exit naïve pluripotency and differentiate. Naïve pluripotent cells have low levels of DNA methylation and instead rely more heavily on H3K9me2/3 for the establishment and/or maintenance of TE silencing^21^. DNA methyltransferases are upregulated as cells exit from naïve pluripotency and contribute to repression of TEs. We noticed that lamin TKO mESCs exhibit moderate activation of TEs, but this activation subsides as TKO cells enter primed pluripotency, which suggests that alternative mechanisms for establishing and/or maintaining TE repression are still functional in lamin-null cells. In contrast, we noted pervasive expression of TEs in both QKO mESCs and EpiLCs. Pluripotent and differentiated cells also differ in their sensitivity to TE expression: differentiated cells express innate immune RNA-sensing proteins that can be activated by TE-derived RNA to induce apoptosis^54–56^ while pluripotent cells do not express these proteins^73^. We noted upregulation of RNA sensors such as *Rig-I* and *Ifit1* in QKO EpiLCs (Fig. 7G), leading us to speculate that TE expression may induce interferon-mediated apoptosis and prevent these cells from surviving the transition to primed pluripotency.

### The nuclear periphery regulates H3K9me2 during differentiation

We have shown that H3K9me2 density increases during the transition from naïve to primed pluripotency. While some earlier reports questioned whether H3K9me2 expands across the genome during differentiation^74,75^, we observed this phenomenon with both immunofluorescence and spike-in-controlled CUT & RUN (Fig. 5J; Fig. 6A-C). Further, we demonstrate that the nuclear periphery regulates the density of H3K9me2 deposition during EpiLC differentiation. When H3K9me2 is displaced from the nuclear periphery by the removal of the lamins and LBR, its density increases to even higher levels in QKO EpiLCs (Fig. 6F-H) but is more disorganized across the genome than in normal EpiLCs (Fig. 6I-J). The levels of the G9a/GLP methyltransferase are known to increase during EpiLC differentiation^24^, but we do not observe further transcriptional upregulation of G9a or GLP in QKO EpiLCs that could explain the abnormally high levels of H3K9me2 observed. We cannot rule out the possibility that the activity of the G9a/GLP enzyme is increased in the absence of the lamins and LBR. We conclude that heterochromatin tethering influences the deposition of H3K9me2 and/or its perdurance on modified loci in primed pluripotency.

Many loci that gain H3K9me2 in EpiLCs are actively transcribed (Fig. 6D-E), which suggests that H3K9me2 performs a distinct function during this developmental transition that is not obligately linked to repression. While the function of H3K9me2 at this stage is poorly understood, recent work indicates that in the epiblast, H3K9me2 expands into actively transcribed regions of the genome that retain chromatin marks associated with active transcription on enhancers and promoters, such as H3K27ac^24,51^. This transitory co-occupation of *cis*-regulatory elements with H3K27ac and H3K9me2 has been proposed to prime these loci for future differentiation-linked silencing^24,51^. The function of H3K9me2 is disrupted in QKO EpiLCs, as H3K9me2 is enriched on dysregulated genes (Fig. 7D). Further, genes associated with various cell fates are discordantly co-expressed (Fig. 7E-H). Therefore, we conclude that recruitment of heterochromatin to the nuclear periphery enables the repression of lineage-irrelevant genes and is required for the normal orchestration of development.

## Supporting information

Merged Supplementary Figures

## Acknowledgements

We are grateful to Joseph Tran and Yixian Zheng for sharing lamin triple knockout and wildtype littermate mESCs, Steve Henikoff for sharing protein-A-MNase for CUT & RUN, Kirstin Meyer and Elphege Nora for guidance culturing and CRISPR-editing mESCs, Karissa Hansen for guidance with RNA-seq library prep and data analysis, the UCSF Genomics CoLab for RNA-seq library preparation and the UCSF CAT core for sequencing of the shLBR dataset, the UCSF Wynton HPC for the computing power that enabled our bioinformatics analysis, John Perrino and the Stanford CSIF for TEM sample preparation and imaging. We thank Geeta Narlikar, Orion Weiner, as well as members of the Buchwalter lab and Nora lab for their helpful discussions since the origins of this project.

A.B. was supported by grants from the National Institute of General Medical Sciences (R35GM142897) and the Chan Zuckerberg Biohub. B. A.-S. was supported by grants from the National Institute of General Medical Sciences (R35GM141888) and National Science Foundation (BIO directorate, 2113319). The project described was supported, in part, by an NIH S10 award from the Office of Research Infrastructure Programs (1S10OD028536-01), although the work described here is solely the responsibility of the authors and does not represent the official views of the NCRR or the National Institutes of Health.

Data for this study was acquired at the Center for Advanced Light Microscopy-CVRI Microscopy core on microscopes purchased though the UCSF Research Evaluation and Allocation Committee, the Gross Fund, and the Heart Anonymous Fund.

## Author Contributions

H.M. and A.B. conceived of the project. H.M., E.S., and A.B. designed experiments. H.M., E.S., C.A., and A.B. performed experiments. B.P., B.A.-S., and A.B. contributed reagents. H.M. generated all cell lines used in this study. H.M. performed immunofluorescence and image analysis. H.M. and E.S. performed CUT&RUN and analysis. H.M. performed RNA-seq and analysis. H.M. and C.A. performed EpiLC differentiation and cell count analysis. E.M. advised on bioinformatics analysis. H.M. and A.B. analyzed data and wrote the manuscript. B.P., B.A.-S., and A.B. supervised the project. All authors discussed results, reviewed, and edited the manuscript.

## Table Titles and Legends

**Table S1.** Genes in H3K9me2 domains in naïve mESCs.

**Table S2.** Genes differentially expressed in naïve mESCs.

**Table S3.** GO analysis of genes differentially expressed in QKO vs TKO mESCs.

**Table S4.** Genes differentially expressed in LBR vs LUC miR-E TKO mESCs.

**Table S5.** TEtranscripts output for TEs DE in naïve mESCs.

**Table S6.** Genes in constitutive and EpiLC-specific H3K9me2 domains.

**Table S7.** TEtranscripts output for TEs DE in EpiLCs.

**Table S8.** Genes differentially expressed in EpiLCs.

**Table S9.** GO analysis of genes differentially expressed in QKO vs TKO EpiLCs.

## Materials and Methods

### Generation of knockout and miR-E expressing mESCs

Lamin triple knockout (vial “TKO97,” passage 12) and littermate wildtype (vial “28538,” passage 9) mESCs were obtained as a generous gift from the laboratory of Dr. Yixian Zheng^76^. Cell lines tested negative for mycoplasma and they were karyotyped by WiCell. Lamin triple knockout mESCs had an abnormal karyotype with two predominant clonal populations: one having trisomy 8, and the other having trisomy 8 and loss of the Y-chromosome. The population of littermate wildtype cells had a 75% normal ploidy with two aneuploid populations: one with loss of the Y-chromosome, and the other with trisomy 6 and 8 and loss of the Y-chromosome.

Lbr knockout was performed by lipofectamine transfection of the PX458 CRISPR/Cas9 plasmid (Addgene #48138) with one guide targeting exon 2 of Lbr (TCATAATAAAGGGAGCTCCC). Transfections were carried out by lipofectamine 2000 (Invitrogen™ 11668019) for Lbr knockout and by electroporation with the Neon™ Transfection System (Invitrogen™ MPK5000) for Lbr rescue experiments using the manufacturer’s procedures. Cells were allowed to recover for 48hrs, then sorted by GFP fluorescence and seeded on a 10-cm dish at a density of 3,000 cells per dish. mESCs were grown for a week until visible colonies appeared, then colonies were picked by pipetting into 96-well plates for screening and further expansion.

The CRISPR guide for Lbr knockout was designed to overlap a restriction enzyme site SacI near the cutting site of Cas9 such that indels would destroy the site. Genotyping was performed by PCR amplification of gDNA with primers flanking the SacI site, then followed by SacI digestion of the PCR product and running the digest on a 2% agarose gel. Homozygous indels were scored by the presence of one band at 1388bp. Heterozygous indels were scored by the presence of three bands at 1388bp, 938bp, and 450bp. Unedited fragments were scored by the presence of two band at 938bp and 450bp. This strategy yielded 36 homozygous Lbr indel clones out of 78 screened in wildtype mESCs, and 10 homozygous Lbr indel clones out of 85 screened in TKO mESCs. The presence of indels was verified by Sanger sequencing and analysis with CRISP-ID^77^. Lbr protein depletion was validated by Western blot with an LBR antibody (cat. ab232731).

To make mESCs expressing tetracycline-inducible miR-Es targeting Lbr or Luc, puromycin, hygromycin, and neomycin resistance markers were first excised from lamin TKO mESCs by transient expression of Cre recombinase and clonal selection. Cre recombinase was expressed from a plasmid derived from pPGK-Cre-bpA (Addgene #11543) with an SV40polyA sequence instead of the bpA. Then, 97mer oligos were ordered containing miR-E targeting LBR (“LBR634” 5′-TGCTGTTGACAGTGAGCGCCAGATATATAGTTACACAGTATAGTGAAGCCACAGATGTATACTGTGTAACTATATATCTGTTGCCTACTGCCTCGGA-3′) and Renilla Luciferase (5′-TGCTGTTGACAGTGAGCGCAGGAATTATAATGCTTATCTATAGTGAAGCCACAGATGTATAGATAAGCATTATAATTCCTATGCCTACTGCCTCGGA-3′). These were amplified by two miR-E universal primers (fwd 5′-CTTAACCCAACAGAAGGCTCGAGAAGGTATATTGCTGTTGACAGTGAGCG-3′) (rev 5′-ACAAGATAATTGCTCGAATTCTAGCCCCTTGAAGTCCGAGGCAGT AGGCA-3′) and cloned into the XhoI and EcoRI double digested LT3GEPIR lentiviral vector (Addgene #111177). HEK293T cells were transfected with these vectors to produce lentivirus which was then used to transduce lamin TKO mESCs. Cells were selected with puromycin (1µg/mL) for 4 days, then individual clones were selected. Using flow cytometry, GFP signal after doxycycline treatment (1µg/mL) was measured to assess the expression of the RNAi system. Clones with a GFP-positive population greater than 88% were selected for experiments.

### mESC culture and EpiLC differentiation

mESCs were cultured in 5% CO_2_ and 37°C under normoxic conditions. mESCs were grown in serum-and feeder-free 2i + LIF medium (N2B27 basal medium, 3 μM CHIR-99021 (Selleckchem S1263), 1 μM PD0325901 (Selleckchem S1036), 10^3^ U/mL LIF (MilliporeSigma™ ESG1107), 55 μM β-mercaptoethanol (Gibco™ 21985023), and 1X PenStrep (GenClone 25-512).) N2B27 medium was made by mixing equal parts of DMEM:F12 (GenClone 25-503) and Neurobasal medium (Gibco™ 21103049) then adding 1X N-2 (Gibco™ 17502001), 1X B-27 (Gibco™ 17504001) and 2 mM Glutamax (Gibco™ 35050061). Dishes were coated with 0.1% gelatin (MilliporeSigma ES-006-B) for 30 minutes at 37°C before seeding cells. Serum-free 2i + LIF media was replenished every day and cells were passaged every other day, seeding 4×10^5^ cells per 6-well (approximately 4.21 x 10^4^ cells/cm^2^).

Differentiation of mESCs into epiblast-like cells was adapted from a published protocol^78^. Briefly, a 6-well was coated with 1 mL of a 1:200 solution of either Geltrex (Gibco™ A1413202) or Cultrex (Bio-Techne 3445-005-01) in N2B27 at 37°C. After one hour, that solution was aspirated and 2 x 10^5^ mESCs were seeded (approximately 2.11 x 10^4^ cells/cm^2^) with N2B27 media containing a final concentration of 20 ng/mL FGF (Peprotech 100-18B) and 12 ng/mL Activin A (Peprotech 120-14E). This EpiLC media was replenished the next day.

For miR-E timepoints followed by RNA-seq, mESCs were seeded at 4.21 x 10^4^ cells/cm^2^ on dishes pre-coated with 0.1% gelatin. mESCs were replenished with fresh 2i + LIF medium every day. Untreated controls were collected after 2 days. Treated cells received doxycycline at 1µg/mL. For day 4 doxycycline timepoints, mESCs were split at day 2, keeping the same seeding density.

### RNA isolation and library preparation for RNA-seq

Four replicates, each consisting of a single 6-well of cells, were used per condition for all RNAseq experiments. Total RNA was isolated using the RNeasy Plus Mini Kit (QIAGEN 74136). RNA was treated with DNAse using the TURBO DNA-free kit (Invitrogen™ AM1907) prior to prepping libraries. RNA quality was assessed by electrophoresis on a 1% agarose gel and detecting 28S and 18S rRNA bands with ethidium bromide staining (0.6µg/mL). Libraries were prepped using the Illumina Stranded Total RNA Prep with Ribo-Zero Plus kit (Illumina 20040525) following the manufacturer’s protocol with a starting RNA input of 500ng. Libraries were amplified using 11 cycles on the thermocycler. Each library was uniquely indexed with IDT for Illumina– DNA/RNA UD Indexes (Illumina 20026121), then pooled together in equimolar amounts. Library concentration was measured using the Qubit dsDNA HS assay kit (Invitrogen Q32851) and Qubit 4 fluorometer. Library size was assayed using the Agilent Bioanalyzer 2100 with the High sensitivity DNA kit (Agilent 5067-4626). Paired-end sequencing was performed on the pooled library using the Illumina NextSeq2000 platform (read length of 150bp and read depth of approximately 50 million reads per library).

For the LBR depletion timecourse, total RNA was isolated using the RNeasy Plus Mini Kit. The Tecan Genomics’ Universal Plus mRNA prep kit (Tecan Genomics 0520B-A01) was modified for rRNA depletion by replacing the mRNA isolation in the first segment of the protocol with ribo depletion using FastSelect rRNA (Qiagen 334387) as follows: 200ng of total RNA in 10µL of water was mixed with 10µL of Universal Plus 2X Fragmentation Buffer and 1µL FastSelect. The solutions were fragmented and ribo-depleted simultaneously per the FastSelect protocol. 20µL of the resulting fragmented/ribo depleted RNA were prepped per the Universal Plus protocol starting with First Strand Synthesis. The Universal Plus Unique Dual Index Set B was used, and the samples were quality checked using a MiniSeq benchtop sequencer (Illumina). Normalized pools were made using the corrected protein coding read counts of each. All concentrations and library/pool qualities were measured via Fragment Analyzer (Agilent Technologies). Paired-end sequencing was performed on pooled libraries using the Illumina NovaSeq6000 platform at the UCSF CAT core (read length of 100bp and read depth of approximately 30 million reads per library).

### RNA-seq analysis

Raw sequencing reads were trimmed of the first T overhang using cutadapt^79^ (version 2.5) as suggested by Illumina. Trimmed reads were mapped to the GENCODE primary assembly M30 release of the mouse genome (GRCm39) using STAR^80^ (version 2.7.10a): --runMode alignReads --readFilesCommand zcat --clip3pAdapterSeq CTGTCTCTTATA CTGTCTCTTATA --clip3pAdapterMMp 0.1 0.1 --outFilterMultimapNmax 1 --outSAMtype BAM SortedByCoordinate - -twopassMode Basic. Bam files were indexed with SAMtools^81^ (version 1.10). The deepTools2^82^ (version 3.4.3) function bamCoverage was used to generate RPKM normalized coverage track files. The subread featureCounts^83^ package (version 1.6.4) and GENCODE M30 primary annotation file was used to generate a read counts table for all conditions, which was used as an input for DESeq2^84^ (version 1.40.2) to obtain normalized counts and perform differential gene expression analysis following a published pipeline which we adapted for mouse (doi.org/ 10.5281/zenodo.3985046). The subread featureCounts package was used again for obtaining read counts on uniquely mapped transposable elements using a custom annotation file made by the Hammell lab (https://labshare.cshl.edu/shares/mhammelllab/www-data/TEtranscripts/ TE_GTF/GRCm39_GENCODE_rmsk_TE.gtf). For Lbr depletion, reads were not trimmed of the first T overhang, but downstream analysis remained the same.

For multimapping of RNA-seq reads, as recommended by the TEtranscripts^85^ authors, STAR parameters were changed from –outFilterMultimapNmax 1 to the following: – outFilterMultimapNmax 100 --winAnchorMultimapNmax 100. A read counts table was generated using the TEcount command from the TEtranscripts package (version 2.2.3) with the following parameters: --sortByPos --format BAM --mode multi. The GENCODE M30 primary annotation file was used for gene annotations and a custom annotation file from the Hammell lab for transposable elements were used to generate a read counts table. DESeq2 analysis was carried out as before.

GO term analysis was performed with Gprofiler^86^. Redundant terms were filtered with ReVIGO^87^.

### CUT&RUN and library preparation

CUT&RUN was performed as described in Skene et al. 2018^88^. Live cells were harvested with accutase and washed once with PBS prior to starting the procedure. Three replicates containing 200,000 mESCs or EpiLCs each were used per condition plus 2,000 HEK293T cells as a spike-in control. H3K9me2 antibody (abcam 1220) was used at 1:100 dilution in antibody buffer. Primary antibody incubation was performed at 4°C overnight. Rabbit anti-mouse secondary antibody (abcam ab6709) was used at 1:100 dilution and incubated for 4°C for 1 hour. pA-MNase (batch #6 143µg/mL) generously gifted from the Hennikoff lab was used for the cleavage reaction.

DNA was isolated by phenol-chloroform extraction. Purified DNA was resuspended with 40µL of 1mM Trish-HCl pH 8.0 and 0.1mM EDTA. DNA concentration was measured using the Qubit dsDNA HS assay kit (Invitrogen Q32851) and Qubit 4 fluorometer. DNA quality was assayed using the Agilent Bioanalyzer 2100 with the High sensitivity DNA kit. Libraries were prepared using the NEBNext Ultra II DNA Library Prep with Sample Purification Beads (NEB E7103S) following the manufacturer’s protocol with 5ng of starting DNA input. Libraries were amplified using 8 cycles on the thermocycler. Each library was uniquely indexed with NEBNext Multiplex Oligos for Illumina (NEB E6440S), then pooled together in equimolar amounts. Library size was analyzed with the Agilent Bioanalyzer 2100 using the High sensitivity DNA kit. Paired-end sequencing was performed on the pooled library using the Illumina NextSeq2000 platform (read length of 35bp and read depth of approximately 16 million reads per library).

### CUT&RUN analysis

Raw sequencing reads were mapped to the “soft masked” mm39 mouse genome using Bowtie2^89^ (version 2.3.5.1): -X 2000 -N 1 --local --dovetail. SAMtools was used to keep properly paired, primary alignments and filter out unassembled contigs, read duplicates, and reads mapped to mitochondria.

For spike-in control scale factor calculations, raw sequencing reads were mapped to the “soft masked” hg38 human genome. As was done for mm39, Bowtie2 and SAMtools were used for alignment and post-alignment processing, respectively. The SAMtools function flagstat was used to find the number of properly paired reads in hg38 alignments. These were used to calculate a scale factor that was defined by dividing an arbitrary constant number (30,000) by the number of properly paired reads (Zheng Y et al (2020). Protocol.io). The scale factors for each library were used to generate bigWig files of 1kb and 10kb bin, spike-in normalized RPKM signal coverage tracks with the deepTools2 function bamCoverage. Four-state hidden Markov models were implemented on each 10kb bin bigwig file with the pomegranate^90^ (version 0.12) package to call four regions of different H3K9me2 signal 1. background signal 2. low signal 3. high signal and 4. very high artifact signal representative of “blacklist” regions. For each condition, replicate BED files containing low (class 1) and high (class 2) H3K9me2 domains were merged using the pybedtools^91^ (version 0.8.1) function multiinter. For each condition, replicate 1kb bigWig files were averaged using deepTools2 function bigwigAverage. From these files, H3K9me2 density in domains were plotted using deepTools2 functions computeMatrix and plotProfile; Kernel density plots were made in base R (version 4.3.1). The subread featureCounts package was used to generate a read counts table for all conditions. For obtaining read counts in genes, the GENCODE M30 primary annotation file was used; for obtaining read counts in transposable elements, the same custom annotation from the Hammell lab was used (see RNA-seq analysis). DESeq2 was used to normalize the raw counts table by library size.

### Immunoblot assay

Frozen cell pellets were lysed with an aqueous buffer containing 8M Urea, 75mM NaCl, 50mM Tris pH 8.0, and one tablet of cOmplete™, Mini, EDTA-free Protease Inhibitor Cocktail (Roche 11836170001). Lysates were run on 4–15% Mini-PROTEAN® TGX™ Precast Gels (BIO-RAD 4561083) and blots were probed with anti-LBR (1:1000, abcam ab232731) and anti-SOX2 (1:1000, abcam ab97959) primary antibodies in 5% milk TBST. Anti-mouse HRP-conjugated (Rockland 610-1302) and anti-rabbit HRP-conjugated (Rockland 611-1302) secondary antibodies were used at 1:5000. Blots were visualized using Pierce™ ECL Western Blotting Substrate (Thermo Scientific™ 32209).

### Cholesterol quantification assay

mESCs (2 x 10^6^ cells per sample) were lysed in 250 uL buffer containing 25 mM HEPES pH 7.4, 150 mM NaCl, 1% Trixon-X-100, 0.1% SDS, and cOmplete™, Mini, EDTA-free Protease Inhibitor Cocktail (Roche 11836170001). Lysate was clarified by centrifugation at 10,000 rpm for 10 minutes at 4C. To determine cholesterol levels, 10 ul of each clarified lysate was analyzed with the Amplex™ Red Cholesterol Assay Kit (Invitrogen A12216) according to the manufacturer’s instructions.

### Immunofluorescence and Halo tag staining

IBIDI chambers (Ibidi USA 80826) were coated with 150μL of a 1:200 solution of Geltrex or Cultrex in N2B27 for 1 hour at 37°C. To obtain well-isolated cells for optimal imaging, 3 x 10^4^ cells in 250 μL media were seeded into each well and allowed to attach in the tissue culture incubator. After approximately 7 hours, mESCs were washed with PBS then fixed with a fresh solution of 4% formaldehyde (Thermo Scientific™ 28908) in PBS for 5 minutes. Cells were washed then stored at 4°C until staining.

For visualizing Halo-tagged constructs, cells were stained before fixation. Briefly, cells were cultured at 37C 5% CO2 with 2i + LIF medium containing 200nM of Janelia Fluor 549 HaloTag ligand (Promega GA1110). After 30 minutes, the media was replaced for N2B27 media and cells were incubated for 5 minutes. Finally, to remove background staining, N2B27 was replaced for 2i + LIF medium and cells were cultured for 1 hour in the incubator until fixation.

Fixed cells were permeabilized in immunofluorescence buffer containing 0.1% Triton X-100, 0.02% SDS, 10mg/mL BSA in PBS. Primary and secondary antibodies were diluted in immunofluorescence buffer and incubated at room temperature for 2 hours and 1 hour, respectively, with immunofluorescence buffer washes in between antibodies. DNA was stained with Hoechst alongside the secondary antibody incubation. Primary antibodies used were H3K9me2 (1:300, abcam ab1220), H3K9me2 (1:1000, ActiveMotif 39041), LAP2 (1:400, Invitrogen PA5-52519), LBR (1:500, abcam ab232731). Secondary antibodies used were anti-mouse Alexa Fluor 488 (1:1000, Invitrogen A-11029), anti-rabbit Alexa Fluor 488 (1:1000, Invitrogen A-11008), anti-mouse Alexa Fluor 568 (1:1000, Invitrogen A-11004), anti-rabbit Alexa Fluor 568 (1:1000, Invitrogen A-11011).

### Image acquisition and analysis

Confocal images were taken using a Nikon spinning disk confocal microscope with a 60 × 1.4 numerical aperture, oil objective. Images were acquired as 0.3µm step Z-stacks with a range of 20 to 50 steps per cell. Unprocessed 16-bit images were saved as ND2 files using the Nikon Elements 5.02 build 1266 software. FIJI was used for cropping, producing max intensity projections, z-slices and converting images to TIFF files. CellProfiler^92^ was used to quantify signal intensity (“MeasureObjectIntensity” module) and radial intensity distribution (“MeasureObjectIntensityDistribution” module). LAP2β staining was used to segment nuclei.

Signal intensity was quantified from max intensity projection images. Radial intensity distribution of H3K9me2 was quantified from a manually selected z-slice that was in the middle of the nucleus. Output values were processed with RStudio and graphed with Prism.

### Transmission electron microscopy and image analysis

mESCs were seeded at density of approximately 4.21 x 10^4^ cells/cm^2^ on 18mm circular coverslips coated with Geltrex (Gibco™ A1413202) in a 12-well dish. EpiLC differentiation was stopped after 32 hours. Cells were fixed in Karnovsky’s fixative: 2% Glutaraldehyde (EMS Cat# 16000) and 4% Formaldehyde (EMS Cat# 15700) in 0.1M Sodium Cacodylate (EMS Cat# 12300) pH 7.4 for 1 hour, chilled and sent to Stanford’s CSIF on ice. They were then post-fixed in cold 1% Osmium tetroxide (EMS Cat# 19100) in water and allowed to warm for 2 hours in a hood, washed 3X with ultra-filtered water, then en bloc stained 2 hours in 1% Uranyl Acetate at RT. Samples were then dehydrated in a series of ethanol washes for 10 minutes each at RT beginning at 30%, 50%, 70%, 95%, changed to 100% 2X, then Propylene Oxide (PO) for 10 minutes. Samples are infiltrated with EMbed-812 resin (EMS Cat#14120) mixed 1:1, and 2:1 with PO for 2 hours each. The samples are then placed into EMbed-812 for 2 hours opened then placed into flat molds with labels and fresh resin and placed into 65C oven overnight.

Cells of interest were located using the grid pattern and cut out with a gem saw and remounted on pre-labeled resin blocks with fresh resin and polymerized overnight again. Once full polymerized the glass coverslip is etched away using hydrofluoric acid for 20 minutes. Using the finder grid pattern left behind the block faces were trimmed down allowing for serial sectioning of the cells of interest.

Sections were taken around 90nm, picked up on formvar/Carbon coated slot Cu grids, stained for 40 seconds in 3.5% Uranyl Acetate in 50% Acetone followed by staining in 0.2% Lead Citrate for 6 minutes. Observed in the JEOL JEM-1400 120kV and photos were taken using a Gatan Orius 2k X 2k digital camera.

Electron signal from TEM images were quantified using FIJI^93^. Forty coordinates were each manually selected for regions beneath the nuclear envelope and the nucleoplasm. Using a FIJI macro, a 0.2µm x 0.2 µm square was drawn centered at each coordinate point, and the integrated signal density was measured. Data from the 40 squares were averaged and used to generate the plot for nuclear envelope to nucleoplasm signal ratio.

## Data availability

Genomic and transcriptomic data generated during the course of this study were uploaded to the GEO database under the following accession numbers: mESC and EpiLC RNAseq data, GSE264599; LBR miR-E RNAseq data, GSE264602; and H3K9me2 Cut & Run data, GSE264603.

