## Supplementary material for "The nuclear periphery confers repression on H3K9me2-marked genes and transposons to shape cell fate": Merged Supplementary Figures

**Supplemental Figure 1**

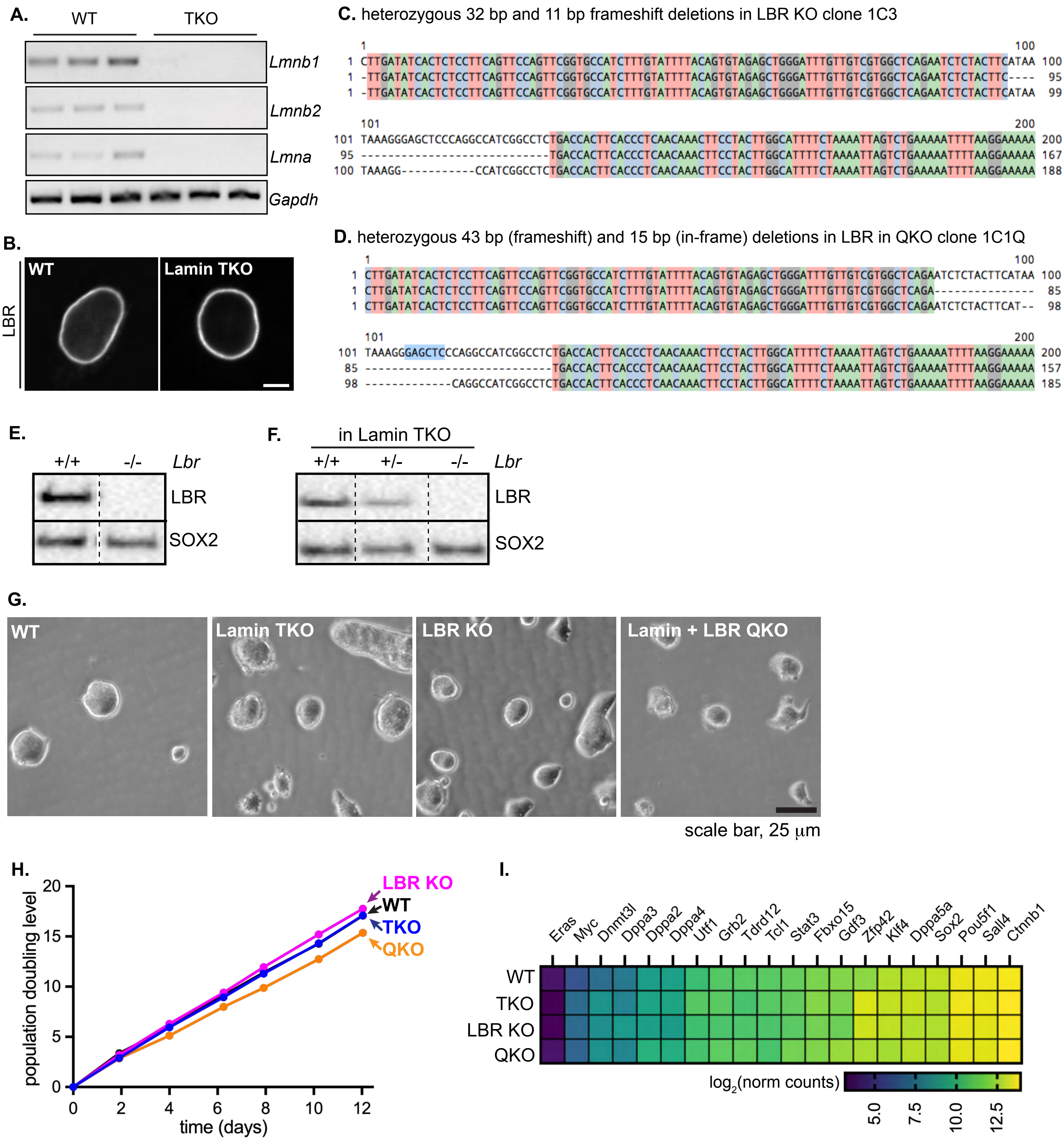

**Supplemental Figure 1.** Validation of knockout mESCs

(A) PCR validation of *Lmna*, *Lmnb1*, and *Lmnb2* knockout in lamin TKO mESCs. (B) Immunofluorescence of LBR in WT and lamin TKO mESCs. Scale bar, 5 mm. Sanger sequencing and CRISP-ID tool analysis showing frameshift indels in LBR KO clone 1C3 (C) and lamin + LBR QKO clone 1C1Q (D). Western blot validation of LBR KO in WT ESC background (E) and in lamin TKO background (F). 10 μg of total protein lysate was loaded for all samples. (G) Colony morphology of WT, lamin TKO, LBR KO, and lamin + LBR QKO mESCs. Scale bar, 25 μm. (H) Growth rate analysis of WT, LBR KO, lamin TKO, and lamin + LBR QKO mESCs. (I) Expression of core pluripotency genes.

Supplemental Figure 2

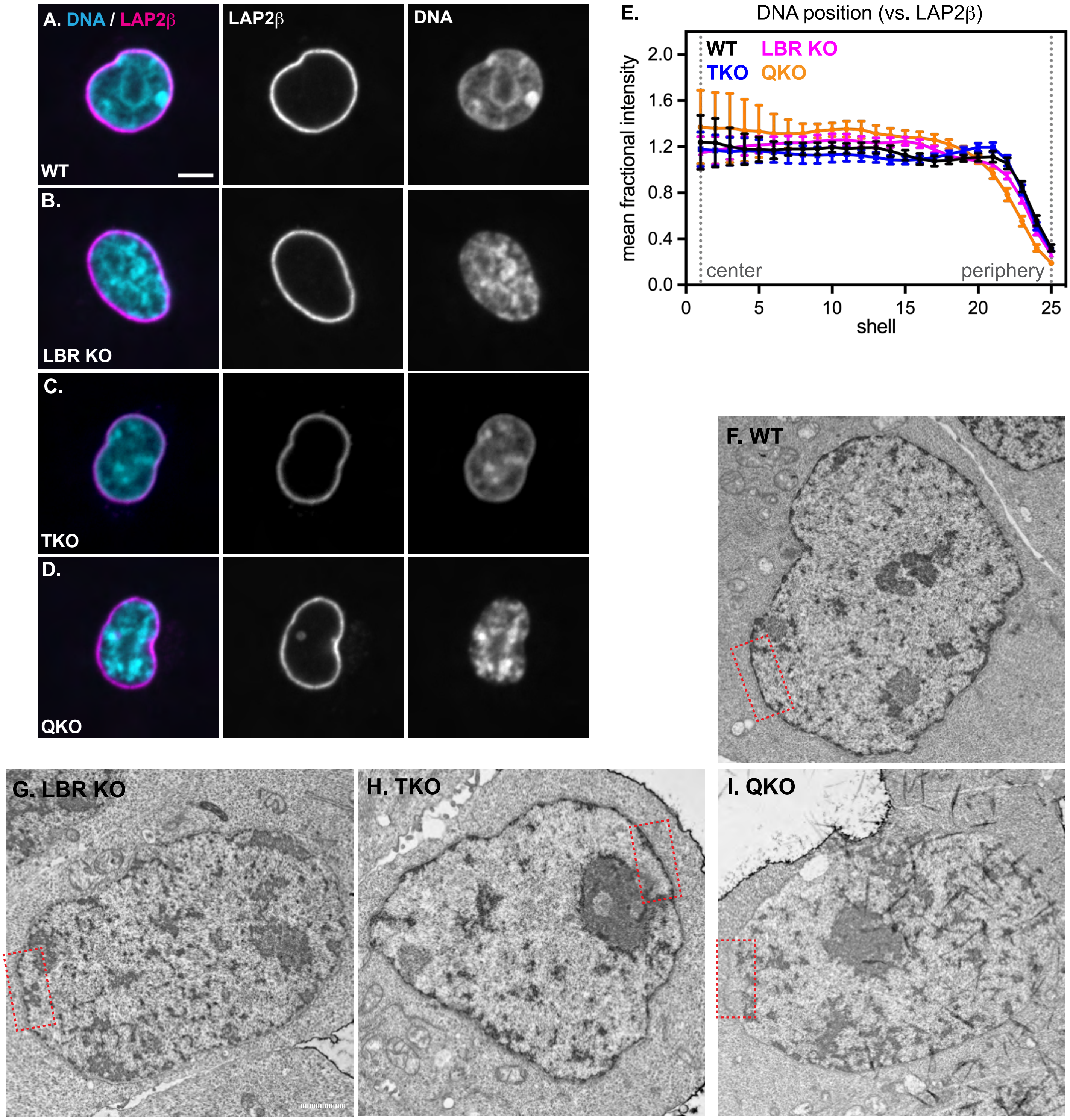

**Supplemental Figure 2.** Microscopy of knockout mESCs.

Immunofluorescence of DNA localization (Hoechst stain) compared to the INM protein (LAP2$\beta$) in WT (A), LBR KO (B), lamin TKO (C), and lamin + LBR QKO (D) mESCs. Central z-slices (XY) are shown. Scale bar, 5 $\mu$m. (E) Radial intensity analysis of DNA position (Hoechst stain) in WT, TKO, LBR KO, and QKO mESCs. ** $p < 0.01$, WT vs QKO shells 8-19, **** $p < 0.0001$, WT vs QKO shells 21-25; * $p < 0.05$, WT vs LBR KO shells 14-18, ** $p < 0.01$, WT vs LBR KO shells 21-25; * $p < 0.05$, WT vs TKO shells 18-21. Transmission electron microscopy images showing full fields of view corresponding to Figure 1 for WT (F), LBR KO (G), lamin TKO (H), and lamin + LBR QKO (I) mESCs. Inset positions that appear in Figure 1 are shown in red dashed boxes.

**Supplemental Figure 3**

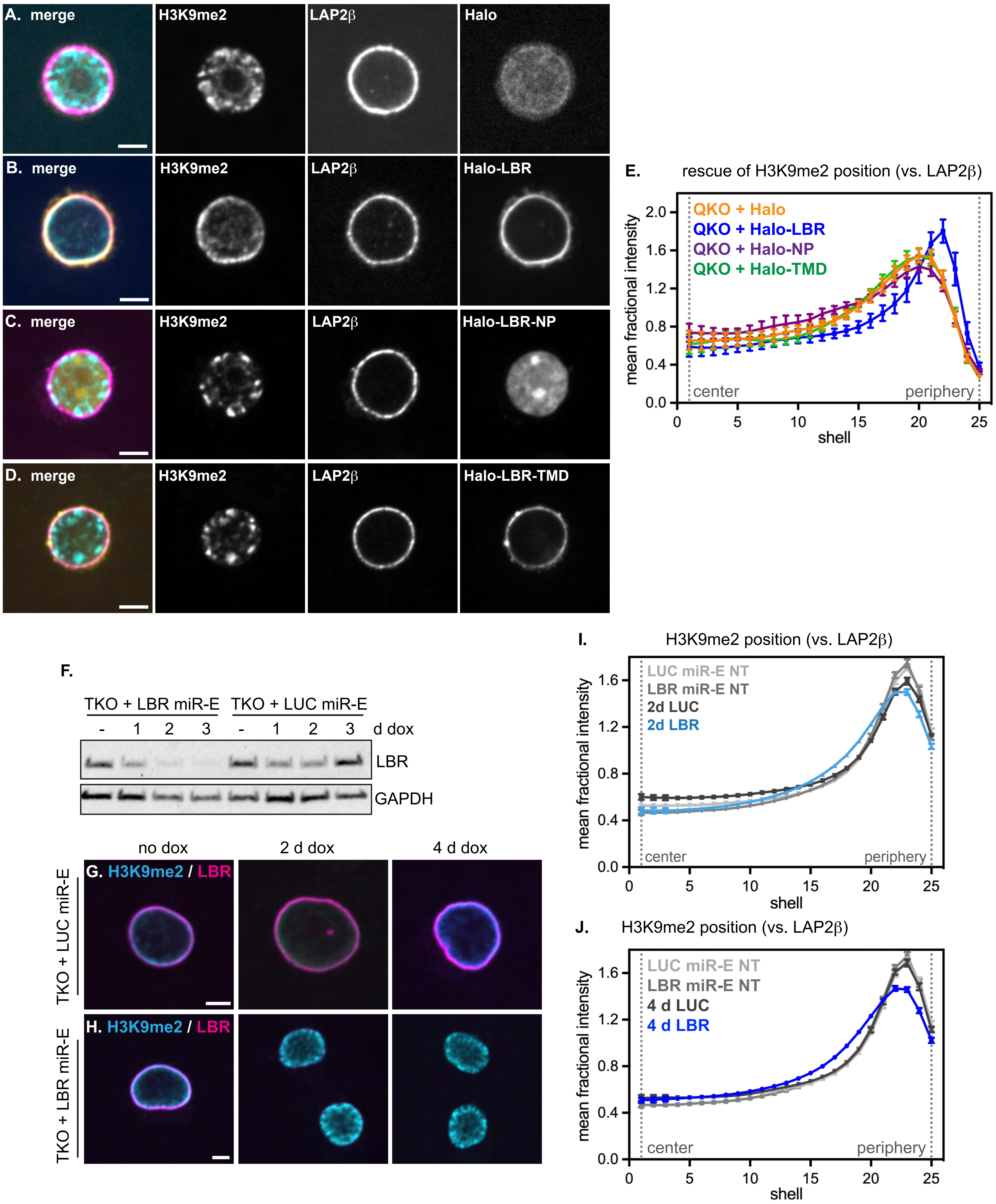

**Supplemental Figure 3.** Displacement of H3K9me2 from the nuclear periphery is reversible.

Immunofluorescence of H3K9me2 localization compared to the INM protein (LAP2$\beta$) in lamin + LBR QKO mESCS expressing Halo-NLS (A), Halo-LBR (B), Halo-LBR nucleoplasmic domain (NP) (C), and Halo-LBR transmembrane domain (TMD) (D). Central z-slices (XY) shown. Scale bar, 5 $\mu$m. (E) Radial intensity analysis of H3K9me2 position in QKO + Halo-NLS (n=27), QKO + Halo-LBR (n=16), QKO + Halo-TMD (n=17), and QKO + Halo-NP (n=26) 2 days after electroporation of plasmids. ** $p < 0.05$, Halo vs. Halo-LBR, shells 12-19; **** $p < 0.0001$, Halo vs. Halo-LBR, shells 22-25, unpaired t-test. ns, $p > 0.05$, Halo vs. Halo TMD and Halo vs. Halo NP in all shells. Points indicate mean and error bars indicate 95% confidence intervals. (F) Western blot showing LBR knockdown in lamin TKO mESCs expressing doxycycline-inducible LBR miR-E. Immunofluorescence of H3K9me2 (cyan) and LBR (magenta) in lamin TKO mESCs expressing LUC miR-E (G) or LBR miR-E (H). Scale bar, 5 $\mu$m. (I) Radial intensity analysis of H3K9me2 position in TKO + LUC miR-E untreated (NT) (n=107), TKO + LUC miR-E + dox 2d (n=140), TKO + LBR miR-E NT (n=102), TKO + LBR miR-E + dox 2d (n=83), unpaired t-test. **** $p < 0.0001$, 2d LBR vs 2d LUC miR-E, shells 1-13, 16-21, 23-25. **** $p < 0.0001$, LBR NT vs 2d LBR miR-E, shells 11-20, 22-24, unpaired t-test. Points indicate mean and error bars indicate 95% confidence intervals. (J) Radial intensity analysis of H3K9me2 position in TKO + LUC miR-E NT (n=107), TKO + LUC miR-E + dox 4d (n=97), TKO + LBR miR-E NT (n=102), TKO + LBR miR-E + dox 4d (n=95), unpaired t-test. **** $p < 0.0001$, 4d LBR vs 4d LUC miR-E, shells 14-20, 22-25. **** $p < 0.0001$, LBR NT vs 4d LBR miR-E, shells 4-20, 22-24, unpaired t-test. Points indicate mean and error bars indicate 95% confidence intervals.

Figure S4

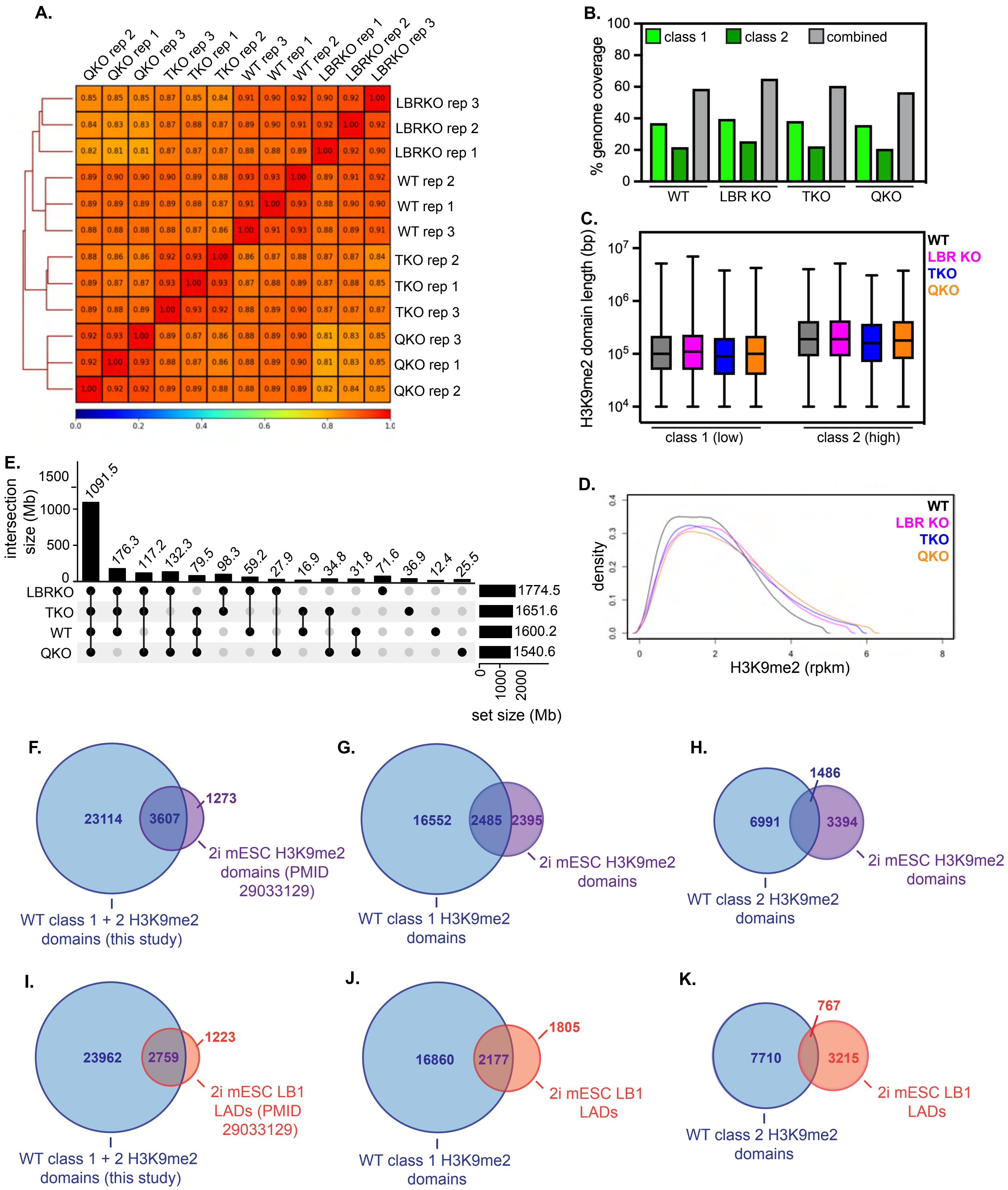


L.

| A vs B | shared (bp) | shared (%) | unique A (bp) | unique A (%) | unique B (bp) | unique B (%) |
|---|---|---|---|---|---|---|
| WT vs LBRKO | 1459423889 | 76.2 | 140753163 | 7.3 | 315112520 | 16.5 |
| WT vs TKO | 1364273204 | 72.3 | 235905471 | 12.5 | 287358524 | 15.2 |
| WT vs QKO | 1335192463 | 73.9 | 264986309 | 14.7 | 205401546 | 11.4 |
| TKO vs QKO | 1323007174 | 70.8 | 328624509 | 17.6 | 217587057 | 11.6 |

**Supplemental Figure 4.** Replicate clustering and analysis of H3K9me2 CUT & RUN in mESCs.

(A) Dendrogram and heatmap of individual H3K9me2 CUT & RUN replicates (3 per condition) showing similarity of replicates for each genotype. (B) Total genome coverage statistics for class 1 H3K9me2 domains, class 2 H3K9me2 domains, and merged H3K9me2 domains in each genotype of mESCs. (C) Genomic length of class 1 and class 2 H3K9me2 domains across genotypes. (D) Histogram of H3K9me2 signal intensity (rpkm) across genotypes. (E) UpSet plot showing all unique overlaps between datasets. (F) Overlap of genes within all merged H3K9me2 domains defined in this study versus in a previous analysis of H3K9me2 in mESCs in 2i + LIF culture conditions (PMID 29033129); 74% of the H3K9me2 genes identified in that study are also found in our dataset. (G) Overlap of genes within class 1 H3K9me2 domains defined in our study versus in PMID 29033129; 51% of the H3K9me2 genes identified in that study are also found in our class 1 H3K9me2 domains. (H) Overlap of genes within class 2 H3K9me2 domains defined in our study versus in PMID 29033129; 30% of the H3K9me2 genes identified in that study are also found in our class 2 H3K9me2 domains. (I) Overlap of genes within all merged H3K9me2 domains defined in our study versus in previously defined LADs (determined by LB1 ChIP-seq) in mESCs in 2i + LIF culture conditions (PMID 29033129); 70% of LB1 LAD genes are also found in H3K9me2 domains in our dataset. (J) Overlap of genes within class 1 H3K9me2 domains defined in our study versus in PMID 29033129; 55% of LB1 LAD genes are also found in our class 1 H3K9me2 domains. (K) Overlap of genes within class 2 H3K9me2 domains defined in our study versus in PMID 29033129; 20% of LB1 LAD genes are also found in our class 2 H3K9me2 domains. (L) Summary of overlapping and unique H3K9me2 domains in pairwise comparisons between genotypes.

**Supplemental Figure 5**

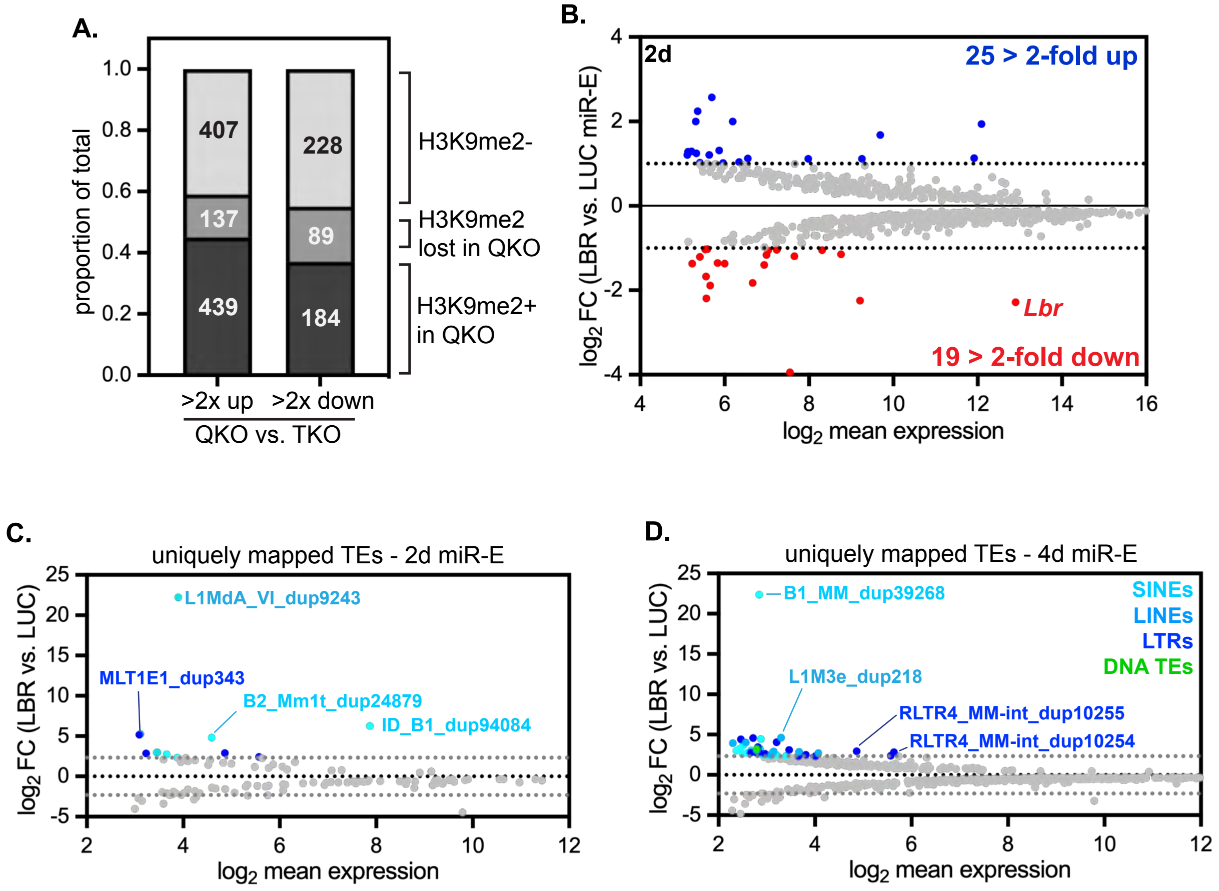

**Supplemental Figure 5.** Analysis of transcription after inducible knockdown of LBR in mESCs.

(A) Analysis of H3K9me2 modification status of genes that are significantly upregulated or downregulated in QKO vs. TKO mESCS, scored as H3K9me2+ (within H3K9me2 domain in QKO), H3K9me2 lost (within H3K9me2 domain in TKO but not QKO) or H3K9me2- (outside of H3K9me2 domains in both genotypes). (B) MA plot comparing gene expression in lamin TKO mESCs expressing LBR or LUC miR-E for 2 days; 778 genes with a minimum $p_{adj}$ of 0.05 shown; 25 genes are upregulated at least 2-fold, while 19 genes are downregulated at least 2-fold. (C-D) MA plots comparing expression of unique TE copies in LBR miR-E versus LUC miR-E after 2 days (C) or 4 days (D). (C) 116 TEs with $p_{adj} < 0.05$ shown; 13 TEs were upregulated at least 5-fold, while 12 TEs were downregulated at least 5 fold. (D) 760 TEs with $p_{adj} < 0.05$ shown; 42 TEs were upregulated at least 5-fold, while 20 TEs were downregulated at least 5-fold. All repeats without a significant change between conditions are gray; significantly differentially expressed TEs (minimum 5-fold change) are colored correspondingly to TE family.

**Supplementary Figure 6**

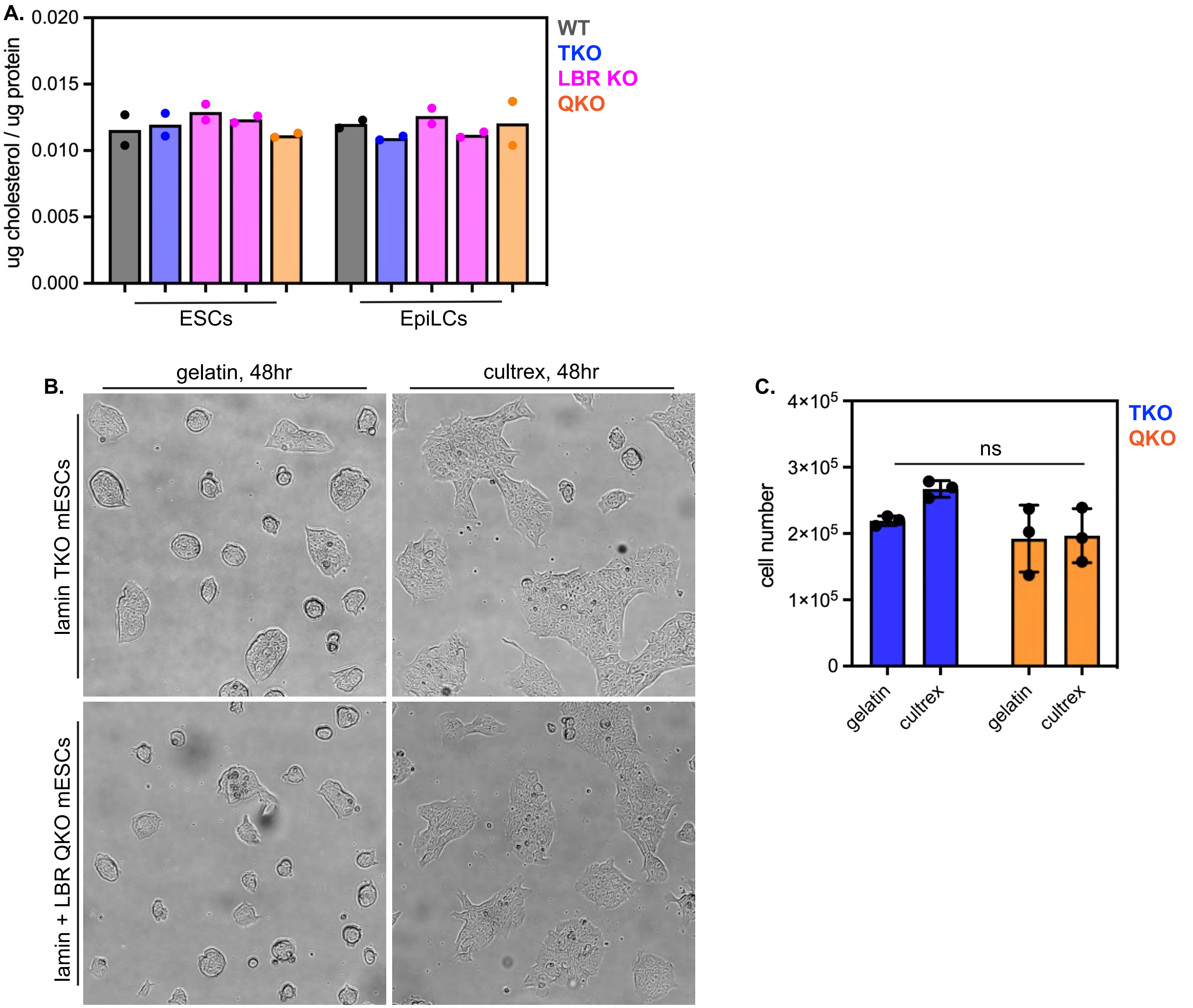

**Supplemental Figure 6.** Additional phenotypic profiling of mESCs.

(A) Representative brightfield microscopy images of lamin TKO and lamin + LBR QKO mESC colonies grown on either gelatin (colonies more rounded with lower attachment) or cultrex (colonies more flattened with higher attachment) for 2 days. (B) Cell numbers of lamin TKO and lamin + LBR QKO mESCs after 2 days of culture on gelatin or cultrex substrate. ns by one-way ANOVA across all conditions; TKO cell numbers are significantly increased on cultrex vs. gelatin ($p < 0.01$). (C) Analysis of cholesterol levels by Amplex Red assay in WT, lamin TKO, LBR KO, and lamin + LBR QKO mESCs and EpiLCs. $n = 2$ replicates per condition.

**Supplemental Figure 7**

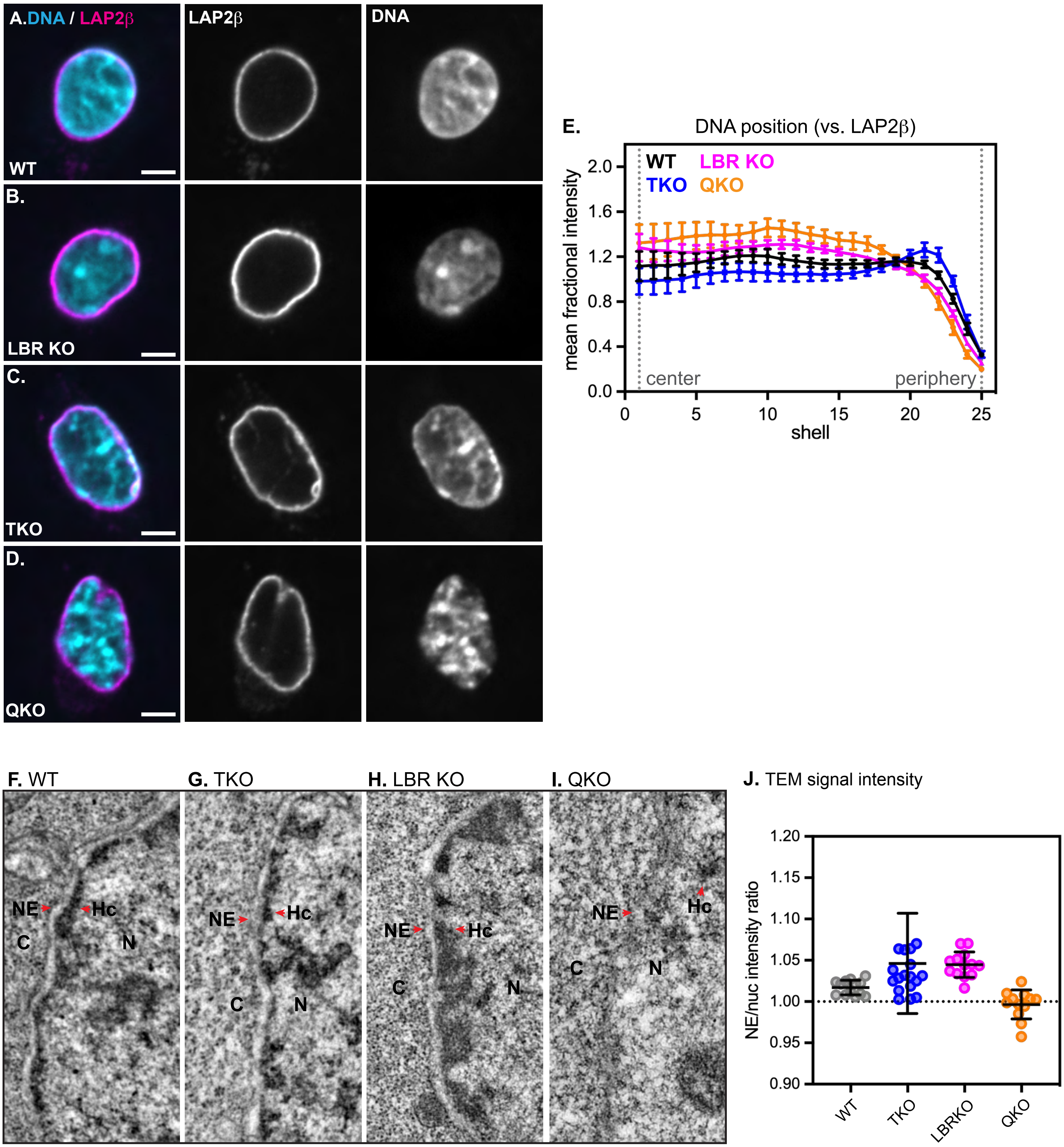

**Supplemental Figure 7.** Fluorescence and electron microscopy of EpiLCs.

Immunofluorescence of DNA localization (Hoechst stain) compared to the INM protein LAP2$\beta$ in WT (A), LBR KO (B), lamin TKO (C), and lamin + LBR QKO (D) EpiLCs. (E) Radial intensity analysis of DNA position (vs. LAP2$\beta$) in WT, TKO, LBR KO, and QKO EpiLCs. ** $p < 0.01$, WT vs QKO shells 5 – 18, **** $p < 0.0001$, WT vs QKO shells 21-25; * $p < 0.05$, WT vs LBR KO shells 8-10, ** $p < 0.01$, WT vs LBR KO shells 11-17, *** $p < 0.001$, WT vs LBR KO shells 20-25; ** $p < 0.01$, WT vs TKO shells 7-16, *** $p < 0.001$, WT vs TKO shells 21-23. Transmission electron microscopy showing 1.3 $\mu$m by 2.6 $\mu$m section of the nuclear periphery in WT (F), lamin TKO (G), LBR KO (H), and lamin + LBR QKO (I) EpiLCs. C, cytoplasm; N, nucleus; NE, nuclear envelope; Hc, heterochromatin. (K) Quantification of relative TEM signal intensity at the NE versus the nucleoplasm for WT (n = 13), LBRKO (n = 18), TKO (n = 13), and QKO (n = 12) EpiLCs.

### Supplemental Figure 8

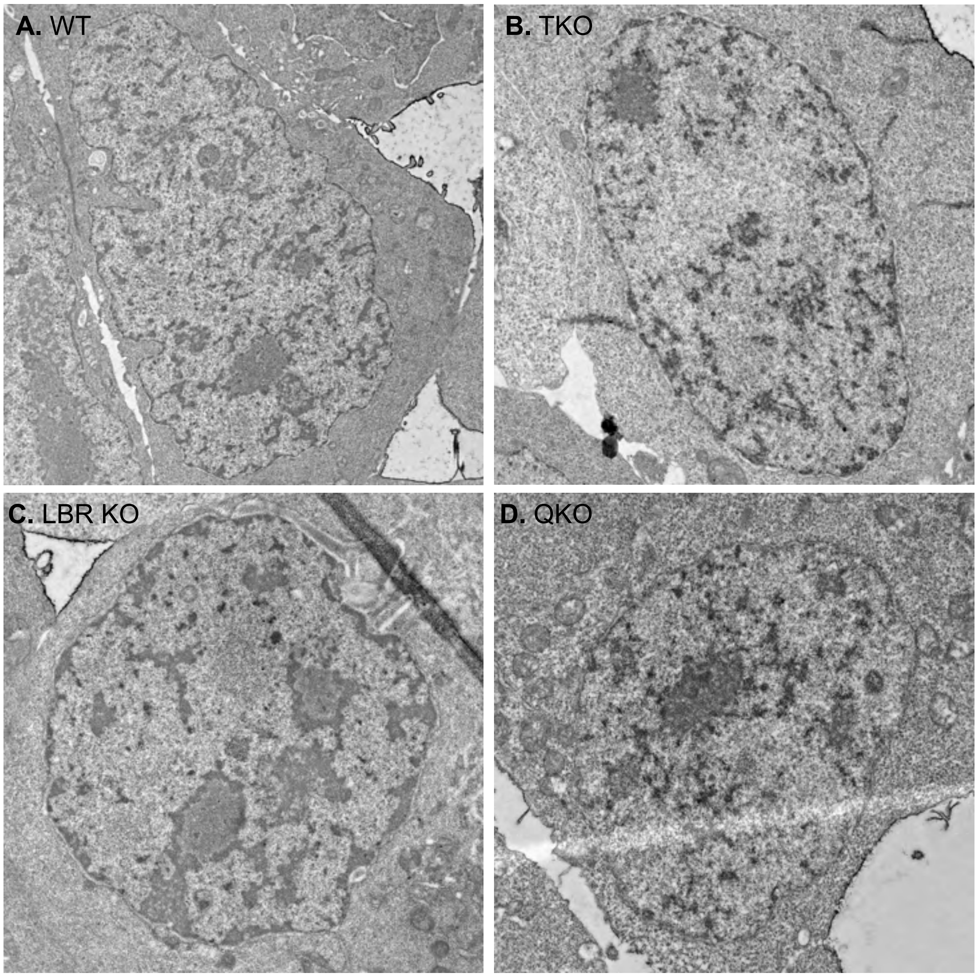

**Supplemental Figure 8.** uncropped TEM images of WT, TKO, LBRKO, and QKO EpiLCs

Figure S9

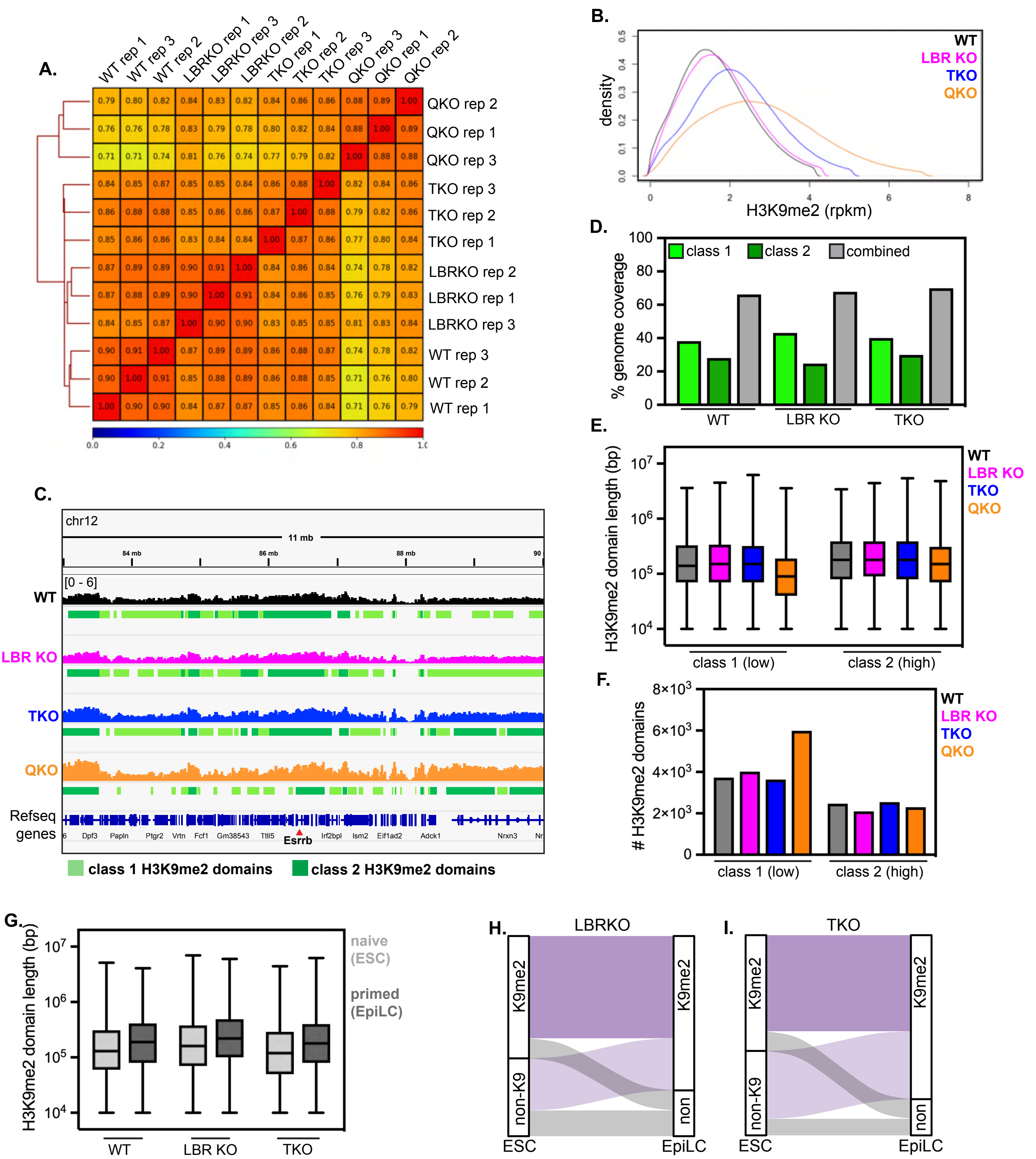


J.

| A vs B | shared (bp) | shared (%) | unique A (bp) | unique A (%) | unique B (bp) | unique B (%) |
|---|---|---|---|---|---|---|
| WT vs LBRKO | 1579407939 | 76.3 | 222751785 | 10.8 | 267217779 | 12.9 |
| WT vs TKO | 1563802588 | 72.9 | 238357179 | 11.1 | 341704668 | 15.9 |
| WT vs QKO | 1249077871 | 58.6 | 553086122 | 25.9 | 329214818 | 15.4 |
| TKO vs QKO | 1321987994 | 61.2 | 583523482 | 27.0 | 256304646 | 11.9 |

**Supplementary Figure 9.** Replicate clustering and analysis of H3K9me2 Cut & Run in EpiLCs

(A) Dendrogram and heatmap of individual H3K9me2 Cut & Run replicates (3 per condition) showing similarity of replicates for each genotype. (B) Histogram of H3K9me2 signal intensity (RPKM) across genotypes. (C) Representative genome tracks and domain calls for H3K9me2 in WT, LBR KO, TKO, and QKO ESCs on a 11 Mb section of chromosome 12, including the *Esrrb* gene. Y-axis range indicated at top left is the same for all tracks shown. Low K9me2 density “class 1” domains are marked in light green and high K9me2 density “class 2” domains are marked in dark green. (D) Total genome coverage statistics for class 1 H3K9me2 domains, class 2 H3K9me2 domains, and merged H3K9me2 domains in each genotype of EpiLCs. (E) Size of class 1 and class 2 domains in each genotype. (F) Number of class 1 and class 2 H3K9me2 domains called in each genotype. (G) Total contiguous merged length of H3K9me2 domains in WT, LBRKO, and TKO EpiLCs versus ESCs. (H-I) Alluvial plots showing movement of genes into and out of H3K9me2 domains as LBRKO (H) and lamin TKO (I) ESCs differentiate into EpiLCs. Genes found in H3K9me2 domains in both ESCs and EpiLCs are referred to as “constitutive” (dark purple) while genes that move into H3K9me2 domains in EpiLCs are referred to as “EpiLC H3K9me2”. (J) Summary of overlapping and unique H3K9me2 domains in pairwise comparisons between genotypes.

Supplemental Figure 10

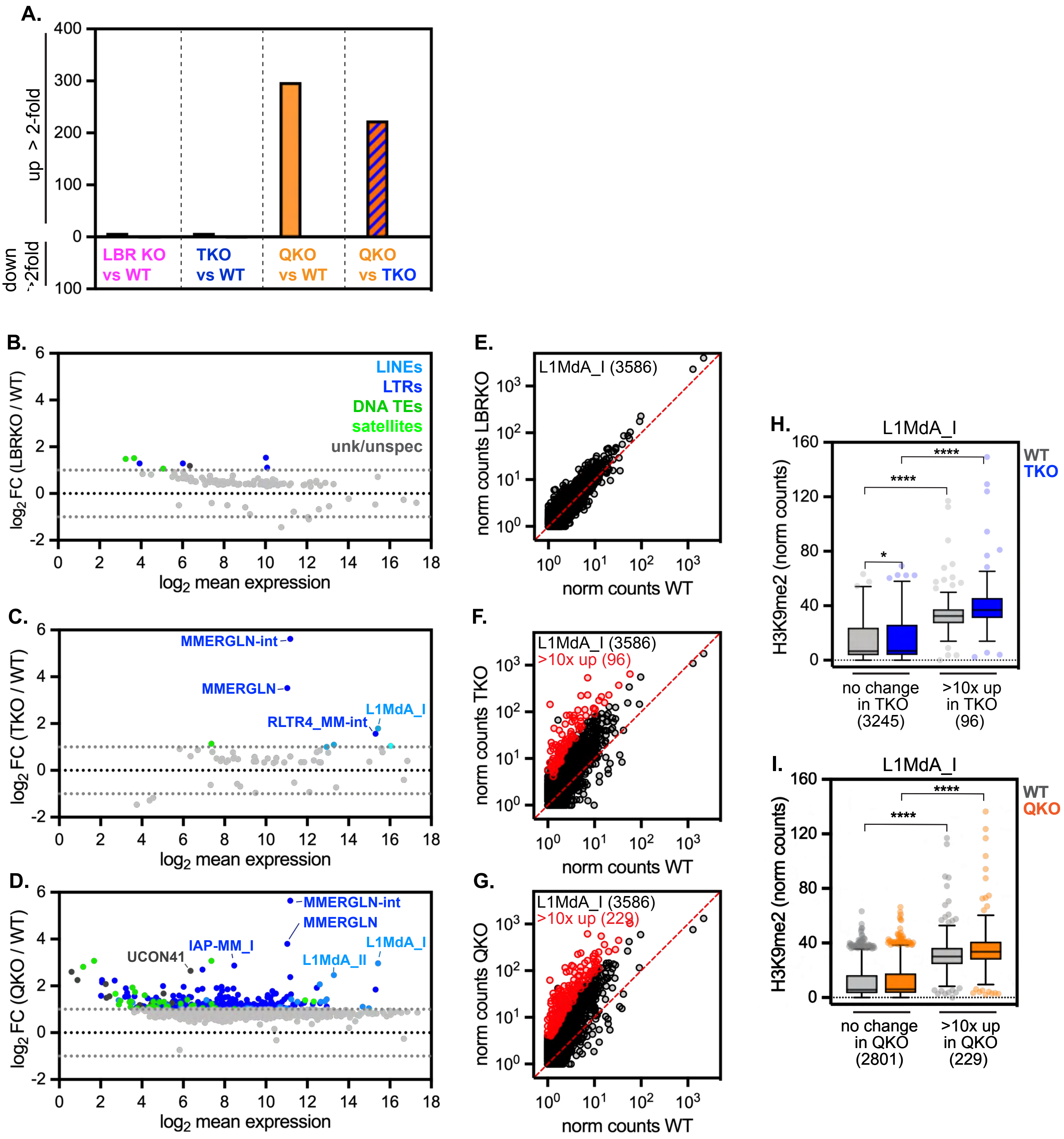

**Supplementary Figure 10.**

(A) Summary of TEs upregulated >2-fold in LBRKO, TKO, and QKO EpiLCs. (B) MA plot of individual TE expression in LBRKO vs. WT EpiLCs. (B) 119 TEs, (C) 56 TEs, and (D) 950 TEs with p adj < 0.05 shown; TEs upregulated at least 2-fold are colored according to TE family. (E-G) Normalized counts for 3586 uniquely mapped L1MdA_I LINE element genomic copies in (E) LBRKO versus WT mESCs, (F) TKO versus WT mESCs, and (G) QKO versus WT mESCs (plotted as $\log_{10}$(average + 1). L1MdA_I copies with >10-fold change and significant difference in expression ($p_{adj}$ < 0.05) in are colored in red. (H-I) Normalized counts from uniquely mapped reads of H3K9me2 on L1MdA_I LINE elements with unchanged expression versus those upregulated >10-fold in TKO mESCs (H, n = 96) and in QKO mESCs (I, n = 229). **** indicates p < 0.0001 and * indicates p = 0.0294 by Kruskal-Wallis multiple comparisons test with Dunn's correction. Box (Tukey) plot center line indicates median; box limits indicate $25^{th}$ to $75^{th}$ percentiles; whiskers indicate 1.5x interquartile range; points indicate outlier values.

.
